# Plasmid co-infection: linking biological mechanisms to ecological and evolutionary dynamics

**DOI:** 10.1101/2021.05.14.444214

**Authors:** Claudia Igler, Jana S. Huisman, Berit Siedentop, Sebastian Bonhoeffer, Sonja Lehtinen

## Abstract

As infectious agents of bacteria and vehicles of horizontal gene transfer, plasmids play a key role in bacterial ecology and evolution. Plasmid dynamics are shaped not only by plasmid-host interactions, but also by ecological interactions between plasmid variants. These interactions are complex: plasmids can co-infect the same host cell and the consequences for the co-resident plasmid can be either beneficial or detrimental. Many of the biological processes that govern plasmid co-infection–from systems to exclude infection by other plasmids to interactions in the regulation of plasmid copy number per cell–are well characterised at a mechanistic level. Modelling plays a central role in translating such mechanistic insights into predictions about plasmid dynamics, and in turn, the impact of these dynamics on bacterial evolution. Theoretical work in evolutionary epidemiology has shown that formulating models of co-infection is not trivial, as some modelling choices can introduce unintended ecological assumptions. Here, we review how the biological processes that govern co-infection can be represented in a mathematical model, discuss potential modelling pitfalls, and analyse this model to provide general insights into how co-infection impacts eco-evolutionary outcomes. In particular, we demonstrate how beneficial and detrimental effects of co-infection give rise to frequency-dependent selection.

## 1 Introduction

Plasmids are mobile genetic elements of bacteria that play a fundamental role in a variety of areas, including bacterial evolution [1, 2], clinical infections [3, 4] and biotechnology [5, 6]. Naturally occurring plasmids exhibit considerable diversity, both in the genes necessary for plasmid replication and spread (’plasmid backbone’) [7–10] - and ‘cargo’ genes, which do not directly impact the plasmid but affect the fitness of the host cell. Such cargo genes can encode traits including antibiotic resistance [11, 12], heavy metal tolerance [13], virulence [14], and toxins for inter-strain competition [15]).

The ecological interactions which shape this diversity are complex: plasmids compete for a limited resource – host cells to infect – but host cells often carry more than one type of plasmid (‘co-infection’) [16–18]. The interactions between co-resident plasmids play a major role in shaping plasmid ecology and evolution. On the one hand, competitive within-cell interactions exert a strong selective pressure on the plasmid backbone, for example by driving the diversification of plasmid replication machinery [19] or the development of systems aimed at hindering co-resident plasmids [8, 10]. Particularly, many plasmids carry systems that prevent co-infection with closely related plasmids, indicating the importance of reducing intra-cellular competition [7]. On the other hand, within-host interactions can also be beneficial for one or both of the co-resident plasmids. This benefit can arise from increased horizontal transmission, for example through increased conjugation rates from co-infected cells to recipient cells [20]; or from vertical transmission (i.e. plasmid inheritance to daughter cells), for example through positive epistasis in fitness cost, meaning that the metabolic burden for the host is reduced [18, 21]. Not all plasmids are conjugative (i.e. can transfer themselves horizontally), but some non-conjugative plasmids can hitchhike along with the conjugation apparatus of co-infecting plasmids [22, 23], making them mobilisable, whereas others are non-mobilisable in general. Overall, within-host interactions crucially shape the fitness landscape plasmids exist in, and thus their population dynamics and diversity.

The (known) biological processes shaping plasmid co-infection have been studied in considerable mechanistic detail [19, 24–27]. Given the complex interactions between these processes and the difficulties in scaling experimental systems to many genetic and environmental conditions, mathematical modelling plays a central role in translating mechanistic insights into predictions about plasmid dynamics and diversity in nature. For example, models of co-infection have provided insights into the conditions for co-existence of conjugative plasmids [28–31]; the maintenance of non-conjugative plasmids [32, 33]; factors influencing gene mobility between plasmids [34]; and the evolution of specific traits such as surface exclusion [28] and toxin-antitoxin systems [35].

Existing models have proven useful in understanding specific aspects co-infection, but here we develop a more general framework relating co-infection processes to eco-evolutionary outcomes. This approach is particularly important because constructing appropriate models of co-infection is not trivial: theoretical work on co-infection between disease strains has shown that seemingly innocuous modelling choices can introduce unintended ecological differences between strains, with considerable impact on model outcomes [36–38]. In particular, model structures easily introduce mechanisms which unintentionally promote strain diversity (‘co-existence for free’) [36]. Models of plasmid conjugation are structurally similar to these epidemiological models of infectious disease transmission, making these concerns about implicit modelling assumptions also relevant for plasmid co-infection.

Our aim is to develop a synthesis of how the biological processes governing co-infection influence the outcomes of plasmid competition. We begin by constructing a general model of co-infection by abstracting many of the processes involved, which allows for flexibility in implementing the underlying biological mechanisms. These different possibilities of implementation are discussed in the context of a literature review on the relevant features of plasmid co-infection. We proceed by giving an intuition of how various co-infection parameters affect bacterial population diversity and by developing a general relationship between co-infection and evolutionary outcomes. Finally, we summarize the main findings of our synthesis and give an outlook on future experimental and theoretical explorations arising from it.

## 2 A model of plasmid co-infection

We begin by developing a model of the population dynamics of two plasmid variants, *A* and *B*, (co-)infecting a bacterial population. This model tracks the density of cell populations in terms of their infection status: no plasmid (*P*_0_), plasmid *A* (*P_A_*), plasmid *B* (*P_B_*) or co-infected with both plasmids (*P_AB_*). We are specifically interested in the effects of vertical and horizontal transmission of co-infection. Hence, our exploration focuses on conjugative plasmids, but the same model structure would also be appropriate for a pair of plasmids where one is conjugative and one mobilisable. The model captures the following fundamental steps in the life-cycle of conjugative plasmids. Plasmids reside within bacterial cells at a copy number determined by the plasmid backbone, which can range from 1-10 [39] to up to 200 [40] copies per cell. (Note that here we do not explicitly model copy number.) Resident plasmids can be transmitted either vertically via host cell replication, or horizontally via conjugation. Vertical transmission requires plasmid replication and partitioning within the cell such that both daughter cells inherit at least one plasmid copy. Conjugation requires expression of transfer genes and close contact between a recipient and a donor cell, allowing transfer of a plasmid copy. The recipient may already carry another plasmid, resulting in co-infection. Co-residence of two (or more) plasmid variants can impact each of these processes and even prevent some from taking place at all. The detailed biological mechanisms will be discussed in section 3. First, we develop a more conceptual intuition of these processes through their realisation in a mathematical model (Figure 1, more details on model structure are given in Supplementary Text 2 and Supplementary Table S1):

**Figure 1:**
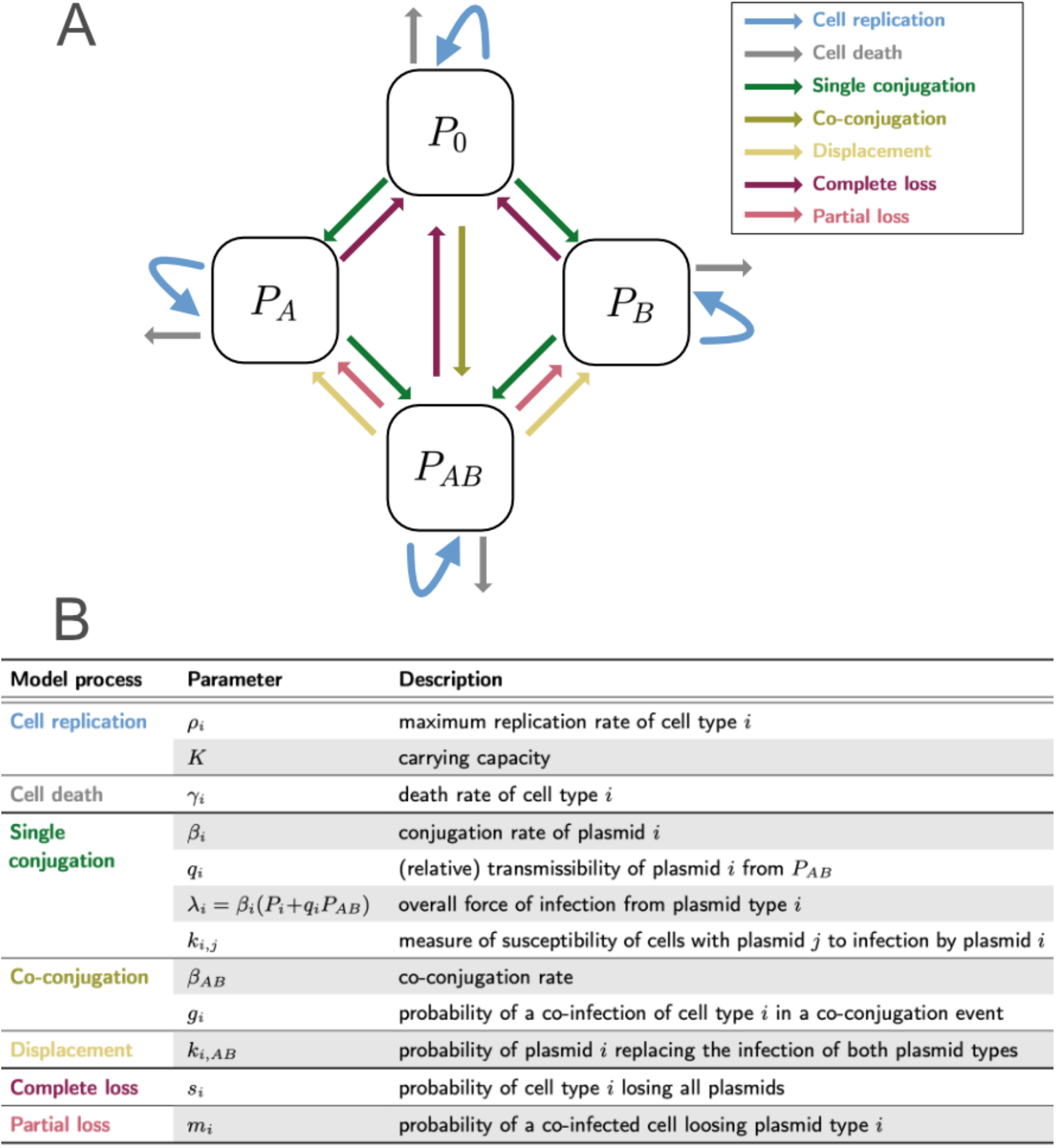
Visualization of the modelled plasmid co-infection processes and the corresponding parameters. **A.** Schematic diagram of the co-infection model given by equations 1. *P*_0_ denotes plasmid-free cells, *P_A_* and *P_B_* are bacterial cells infected with plasmid variant *A* or *B*, respectively, and PAB are cells co-infected with *A* and *B*. Arrows indicate the transition of cells between states. **B.** Co-infection processes incorporated in the model, listed with their associated parameters and parameter descriptions.

### Bacterial population size

We model changes in the host cell density in two components: i) a density-dependent *replication* rate 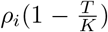, with *ρ_i_* representing the maximum replication rate, *i* the cell type (0, *A, B* or *AB*), *T* the total cell density (*T* = *P*_0_ + *P_A_* + *P_B_* + *P_AB_*) and *K* the carrying capacity; ii) a density-independent *death* rate *γ_i_*. Plasmid costs and benefits can be captured in both *ρ_i_* and *γ_i_*, for each cell type individually.

### Plasmid conjugation

#### Single conjugation

Plasmids conjugate in a manner dependent on host cell density, at rate *β_i_*, where *i* indicates plasmid variant *A* or *B*. The relative transmissibility of plasmid *i* from co-infected cells (*P_AB_*), is given by *q_i_*. Thus, the overall force of infection from plasmid variant *i* is *λ_i_* = *β_i_*(*P_i_* + *q_i_P_AB_*).

If the recipient cell is already (singly) infected with plasmid variant *j*, further infection with plasmid variant *i* is possible, and leads to co-infection. The susceptibility of cells with (only) plasmid *j* to infection by plasmid *i*, relative to cells with no plasmid, is given by *k_i,j_*.

If the recipient is already co-infected, further infection with either variant can theoretically lead to *displacement* of the co-resident variant, and a return to a singly infected state (known as ‘knock-out’ in the epidemiological modelling literature [36]). The probability of plasmid *i* displacing plasmid *j* from a co-infected cell upon infection is given by *k_i,AB_*.

#### Co-conjugation

If co-infected cells can also transmit both plasmids simultaneously (‘co-transfer’), co-conjugation from *P_AB_* occurs at rate *β_AB_*. Hence, the overall infectiousness of co-infected cells is given by *q_A_β_A_* + *q_B_β_B_* + *β_AB_*. If the recipient carries no plasmid (*P*_0_), it transitions directly to the *P_AB_* state. If the recipient is singly infected, e.g. *P_A_*, co-conjugation leads to co-infection with probability *g_A_.*

### Plasmid segregation loss

#### Complete loss

Cells can lose (single or double) plasmid carriage completely during cell division (*s_i_*).

#### Partial loss

Co-infected cells can revert to being singly infected if they lose only one plasmid variant. This occurs with probability *m_i_* (with the constraint *m_A_* + *m_B_* ≤ 1). Note that, depending on the specific mechanism of plasmid loss in co-infected cells, *s_i_* and *m_i_* may not be independent, which can be captured by constraining their relationship.

These processes are captured by the following equations (with colors corresponding to Figure 1):

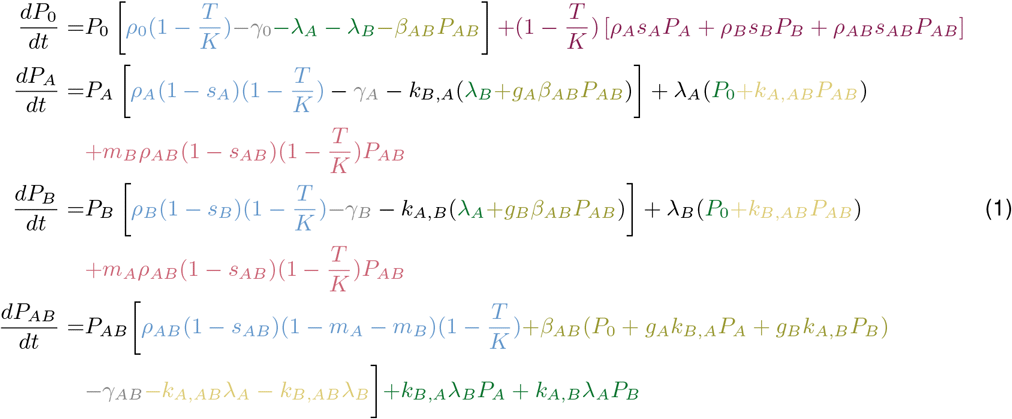

## 3 Model parameters - Biological mechanisms

Having introduced the basic processes involved in plasmid co-infection, we will briefly portray the underlying complexity of biological mechanisms and how these can be incorporated into our model structure.

### Plasmid replication and partitioning

The most important steps in faithful vertical plasmid transmission are plasmid replication and (for some plasmids) partitioning, which positions plasmid copies within the cell to ensure inheritance to both daughter cells. When co-infecting plasmid variants share the same replication and/or partitioning regulation, either variant is more likely to be lost during cell division. This leads to an inability of plasmid variants to coexist stably in the same cell lineage, which is used to define plasmid incompatibility [19] – although, as this definition is based on a phenotype, ‘incompatibility’ can also arise from other within-host interactions [41]. The degree of incompatibility is dependent on the specific system and the plasmid variant, with identical co-resident plasmids showing segregation loss rates between 1-15% per replication [42] (Table S2).

#### Replication systems

Plasmid replication, and hence plasmid copy number in the cell, is tightly regulated to minimize the cost to the host - while still guaranteeing stable vertical transmission. Generally, the distribution around the target copy number within each cell is a narrow Gaussian [43], but recent evidence showed that the standard deviation can be on the order of the mean copy number [44]. Replication control is based on feedback from the plasmid copy number in the cell (down-regulation at high copy numbers)[19]. Hence, incompatibility arises from the inability of plasmids to differentiate between their own and the co-resident’s copy number and correct for deviations from the target number [19]. Two plasmid variants sharing replication determinants will establish the same overall copy number as they would individually, but with a mixed plasmid pool. Random sampling from this pool for replication leads to heterogeneity in the within-host frequencies of the two plasmid variants [19]. In absence of other effects (including conjugation), genetic drift will lead to eventual loss of all copies of one plasmid variant from the cell lineage (Table S2).

#### Partitioning systems

To ensure stable inheritance to both daughter cells, sibling plasmids have to be separated into the two cell halves after replication. This is especially important for low copy number plasmids, which are known to use partitioning systems for this purpose. However, non-random positioning has also been found for high copy number plasmids [45], which is beneficial if heterogeneity in copy number can indeed be large [44].

Partitioning systems generally consist of three (plasmid-encoded) components: a centromerelike DNA site and two proteins, an NTPase (energy production and movement) and a centromere-binding protein (plasmid tethering) [46]. The incompatibility mechanism is determined by the affected component and can lead for example to random partitioning or centromere-binding protein sequestration [47]. The variation that is found in centromere-like DNA sites alone indicates selection pressure for distinct partitioning systems [47]. Notably, some plasmids harbor multiple partitioning systems, which can increase their stability compared to either system alone [48].

The influence of partitioning and replication systems on plasmid co-infection differs depending on their relatedness (Figure S6):

- Identical replication systems: Complete and partial segregation loss are symmetrical (*s_AB_* = *s_A_* = *s_B_*, *m_A_* = *m_B_*). Partial segregation loss is more frequent than for compatible plasmids (Table S2), especially if partitioning is also incompatible [49].
- Related replication systems: Partial segregation probabilities can be either symmetric or favor the plasmid that is less sensitive to the incompatibility determinant. Higher stability could also be related to a difference in copy number, as higher numbers increase the chance of being selected as a replication template [19].
- Compatible replication systems: Incompatibility can still arise via partitioning systems only. Again, this can lead to symmetric or asymmetric segregation loss for co-resident plasmids. Interestingly, for low copy number plasmids with partitioning incompatibility, loss rates can be even higher (4-5fold) than those arising from random partitioning [50].

Replication and partitioning further influence susceptibility to co-infection and displacement (*k_i,j_, k_i,AB_*). First, a newly co-infecting plasmid variant will have a low copy number compared to the established variant, thus making it more likely to be lost during the first rounds of cell replication, if the previously established plasmid is incompatible. Second, if segregation loss of one of the incompatible plasmid variants is very rapid, co-infection becomes negligible and need not be modelled at all. Current estimates indicate however, that plasmid loss is slow, with probabilities of 1-22% per generation (Table S2).

Replication and partitioning systems impact a number of other model parameters indirectly, since they lead to a lower copy number of each plasmid variant in the co-infected cell. This can decrease the probability of successful conjugation (*q_i_*) [51] and plasmid cost (*ρ_AB_, γ_AB_*), compared to co-infection with compatible plasmids.

### Toxin-Antitoxin systems

Toxin-Antitoxin (TA) systems on plasmid are usually seen as addiction modules that select against plasmid-free cells through ‘post-segregational killing’ [52]: After plasmid loss, neither toxin nor antitoxin is produced any longer, but the more stable toxin persists (without antitoxin) in the cell and interferes with essential cellular processes like replication, translation and cell-wall synthesis [53]. However, toxin inhibition of cell metabolism seems generally reversible (e.g. the F plasmid toxin inhibits cell division only until completion of plasmid replication [54]), with cell killing only being observed in over-expression experiments [55]. This suggests that TA systems not only reduce competition from cells that have lost the plasmid, but also increase faithful inheritance by slowing cell division.

While TA systems have been found to promote plasmid maintenance, they seem to be (up to a 100-fold) less efficient than partitioning systems [53] (Table S2). Their overall stabilization effect varies considerably (2.5-100fold) and is dependent on the host strain [56] (Table S2). The impact of TA systems during co-infection could be greater, as loss of the TA-carrying plasmid will slow down vertical and horizontal transmission of the non-TA-carrying plasmid [8].

The influence of plasmid TA systems can be modeled in various ways (Table 1):

- If TA systems kill the plasmid-free host, segregation loss leads to cell death instead of transition to the plasmid-free state. This can be modelled by introducing a (1 – *x*) modifier to the complete segregation loss term (*s_i_*) in the equation for *P*_0_ (only): a proportion *x* of cells that lose the plasmid die. For co-infection with a TA-carrying (*A*) and non-TA-carrying (*B*) plasmid, partial segregation loss (*m_A_*) and displacement (*k_B,AB_*) can be similarly modified in the equation for PB to capture cell death following the loss of plasmid *A*.
- If TA systems slow down cell division, the increased vertical stability can be modelled by decreasing complete (*s_i_*) and partial segregation loss (*m_i_*), at the cost of a lower replication rate (*ρ_i_*). This slower cell division may also increase vertical stability (i.e. decrease *m_i_*) of a co-resident plasmid. The decreased competitiveness of cells that have lost the TA-carrying plasmid would be most accurately represented by introducing additional states to capture the temporary reduction in post-segregational replication rate. To avoid the introduction of additional states, the effect may be approximated by modelling the decreased net growth rate through post-segregational death (i.e. as above).

**Table 1:**
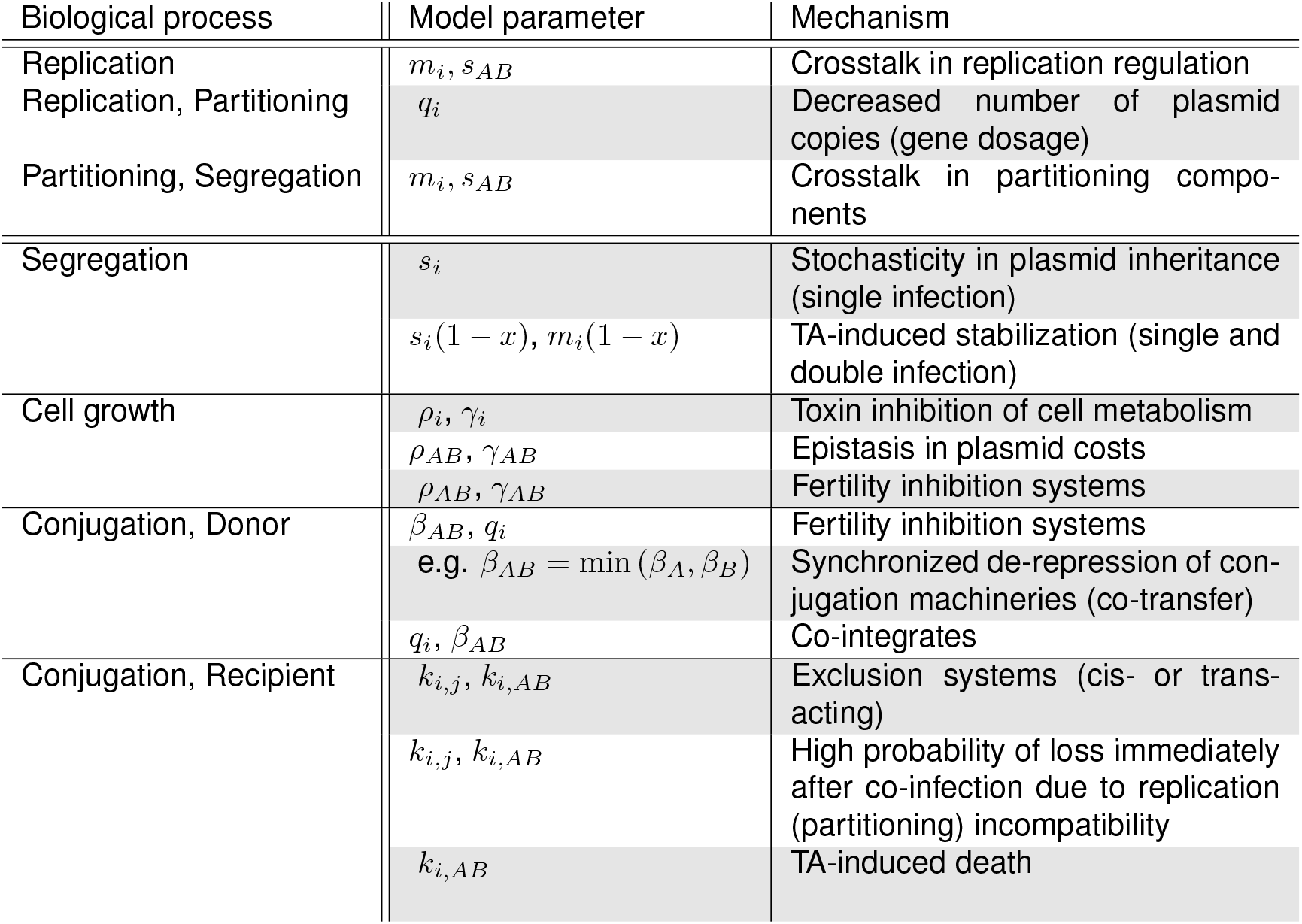
Summary of biological processes relating to co-infection and their relation to model parameters.

| Biological process | Model parameter | Mechanism |
| --- | --- | --- |
| Replication | $m_i, s_{AB}$ | Crosstalk in replication regulation |
| Replication, Partitioning | $q_i$ | Decreased number of plasmid copies (gene dosage) |
| Partitioning, Segregation | $m_i, s_{AB}$ | Crosstalk in partitioning components |
| Segregation | $s_i$ | Stochasticity in plasmid inheritance (single infection) |
| | $s_i(1 - x), m_i(1 - x)$ | TA-induced stabilization (single and double infection) |
| Cell growth | $\rho_i, \gamma_i$ | Toxin inhibition of cell metabolism |
| | $\rho_{AB}, \gamma_{AB}$ | Epistasis in plasmid costs |
| | $\rho_{AB}, \gamma_{AB}$ | Fertility inhibition systems |
| Conjugation, Donor | $\beta_{AB}, q_i$ | Fertility inhibition systems |
| | e.g. $\beta_{AB} = \min(\beta_A, \beta_B)$ | Synchronized de-repression of conjugation machineries (co-transfer) |
| | $q_i, \beta_{AB}$ | Co-integrates |
| Conjugation, Recipient | $k_{i,j}, k_{i,AB}$ | Exclusion systems (cis- or trans-acting) |
| | $k_{i,j}, k_{i,AB}$ | High probability of loss immediately after co-infection due to replication (partitioning) incompatibility |
| | $k_{i,AB}$ | TA-induced death |

### Effect on host cell fitness

The effect of plasmids on the fitness of their host cells can be negative or positive. Hence, co-infection can impact the vertical transmission of co-resident plasmids through effects on host cell replication or death (*ρ_AB_, γ_AB_*). Importantly, these effects may be different than expected from the effects of each plasmid individually (epistasis). For example, there is empirical evidence of positive epistasis (i.e. reduced fitness costs) between co-infecting plasmids [18, 21], which could stem from down-regulation of the conjugation machinery [57] (see below) and/or a decrease in the number of individual plasmid copies per cell [58]. Epistatic effects could also arise from interactions between plasmid cargo genes (e.g. resistance to the same antibiotic).

### Conjugation from co-infected cells

A key characteristic of conjugative plasmids is their ability to transmit themselves horizontally to neighbouring cells, which requires the expression of transfer genes from the plasmid, and close proximity between the recipient and donor cell.

To reduce the burden on the host, the conjugation machinery is generally down-regulated (‘repressed’) and not active at all times [59]. Plasmids typically carry fertility inhibition (FI) systems, which inhibit conjugation, either as an auto-regulatory mechanism (F plasmids), or to inhibit transfer of unrelated, co-resident plasmids [10, 60] (Table S2). Activation is also influenced by diverse factors such as host cell physiology, the availability of recipients, or stress factors like antibiotics [61, 62]. Such external activation signals can de-repress both co-infecting plasmids, increasing the chance of simultaneous transfer [63].

Co-infecting plasmids can affect each other’s individual conjugation rates (*q_A_,q_B_*), as well as transfer simultaneously during a single mating event (co-transfer; *β_AB_*). Effects on individual conjugation rates during co-infection are common (63% of tested plasmid pairs), although typically only one plasmid is influenced (53% of plasmid pairs) [20]. In this case, a reduction in conjugation rate is more common (30%) than an increase (23%) [20].

Co-transfer of plasmids can occur through the same type IV secretion system (T4SS), or by expression of several systems simultaneously. Mobilisable plasmids can ‘hitch-hike’ along with the T4SS of a conjugative plasmid, if they encode compatible transfer determinants [22, 23]. Transfer via the same T4SS can also occur with plasmid co-integrates [64], which arise through fusion of plasmid variants.

In the case of multiple co-resident, conjugative plasmids, simultaneous expression of secretion systems could stabilise the mating pair, thus allowing efficient co-transfer [20]. However, determination of the true rate of conjugative co-transfer is difficult as ‘simply’ counting the number of recipients that received both plasmids makes it difficult to distinguish whether a single or two subsequent mating events took place. This may explain the variation in empirical co-transfer reports, showing frequent co-transfer in a system with large and small plasmids [65], and in an engineered system with conjugative plasmids [63], but little in another system with conjugative plasmids from natural isolates [66].

The effect of co-infection on conjugation can be modelled in the following ways (Table 1):

- FI systems decrease the single and co-conjugation rate (*q_i_, β_AB_*) of co-resident plasmids, resulting in up to 10,000-fold lower conjugation rates [60]. Lower conjugation rates might in turn decrease the plasmid burden on the host cell (*γ_AB_, ρ_AB_*) [57].
- Co-transfer rates of co-resident plasmids are largely unknown, but have been proposed to occur at the rate set by the lower conjugation frequency (*β_AB_* = min (*β_A_, β_B_*)) [63].
- Co-integrates, i.e. fused plasmid variants, can increase (higher probability of expressing at least one conjugation machinery) [67] or decrease (lower mating pair stability) the rate of co-conjugation (*β_AB_*), and hence the total conjugation frequency of individual plasmids (*q_i_β_i_* + *β_AB_* ≶ *β_i_*). Note that our model only captures this process if co-integrates are resolved again.

### Cis-acting prevention of co-infection

Conjugative plasmids carry genes with which they can prevent co-infection by plasmids from the same exclusion class (i.e. cis-acting) [7]. This serves to reduce i) within-host competition between plasmids, ii) the metabolic burden of conjugation on donor cells, and iii) recipient death due to excessive DNA transfer (lethal zygosis) [7]. There are two types of exclusion systems: surface exclusion (SFX), which inhibits the ability to form stable mating pairs, and entry exclusion (EEX), which inhibits DNA transfer across the mating channel. While the latter is found in nearly all conjugative plasmids, only plasmids with pili that firmly attach to the recipient cell code for surface exclusion [7, 60].

For F plasmids, entry exclusion was found to be around 10 times more effective than surface exclusion [9, 25, 26, 68]. Together, these systems can generate differences in plasmid transfer between 100-10’000-fold (individually, 200- and 20-fold for EEX and SFX, respectively) [25, 26, 68]. Similarly, 10-10’000-fold reductions in transfer have been observed for EEX with other incompatibility groups [7, 69]. The width of this range is likely due to differences in plasmid copy number, as exclusion was found to be gene dosage dependent [7, 68, 69].

Despite the ubiquity of exclusion systems, in practice their impact remains unclear. First, there is substantial genetic diversity between SFX and EEX genes, and how this translates into the exclusion phenotype is not well understood. Within the group of F-like plasmids, at least four different surface exclusion groups were identified [70], where specificity was determined only by a difference of 5 amino acids [71]. The EEX gene is less conserved than the SFX gene: homologous EEX genes were found in only 30% of 256 F-plasmids [72]. Second, certain broad host range plasmids exhibit ‘retrotransfer’, whereby the plasmid is transferred into a recipient, ‘captures’ chromosomal genes or a mobilisable plasmid from that recipient, and is then able to transfer back into the original plasmid-carrying host [73]. Third, little is known about the effect co-resident plasmids have on exclusion. In one experiment, a donor with two plasmids carrying different SFX systems managed to infect a recipient with either one of these plasmids [70]. Fourth, plasmids can bypass exclusion systems by being taken up via a different route (e.g. via transformation, transduction or vessication) [1]. Lastly, exclusion is not active when recipients are in stationary phase [70, 74], allowing infection by plasmids from metabolically active donors, or by plasmids that can conjugate in stationary phase [61].

In our model, the parameters describing co-infection susceptibility *k_i,j_* and displacement *k_i,AB_* can account for exclusion (Table 1):

- If exclusion systems are highly effective, modelling co-infection is only relevant for plasmids of different exclusion groups. Co-infected cells would exclude further entry and displacement by either plasmid type (*k_i,AB_* = 0).
- With less effective exclusion systems, cells may be infected by plasmids of the same exclusion group. Displacement *k_i,AB_* is thus greater than 0, independent of the exclusion group of plasmid *A* and *B*. If co-infecting plasmids are of the same exclusion and incompatibility group, then *k_i,AB_* is constrained to 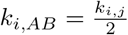 to retain structural neutrality (see Supplementary Text 3) [36].

### Trans-acting prevention of co-infection

Plasmids can also affect the entry and establishment of other variants into a cell in trans, for example via restriction modification (RM) systems and CRISPR (clustered regularly interspaced short palindromic repeats) plus associated Cas genes (CRISPR associated systems) [60, 75, 76].

Restriction-modification systems consist of two functional parts: one cleaves DNA at specific restriction sites, and the other continuously modifies (methylates) these sites to avoid cleavage. This serves primarily as defence against incoming, non-methylated DNA, which will be cleaved upon entry. DNA within the same cell is protected, as long as methylation is actively maintained. If an RM system is lost and the methylation wears off, the remaining restriction endonucleases can kill the cell (i.e. akin to post-segregational killing by TA systems). RM systems are typically located on the chromosome, but are also found in approximately 20% of mobilisable and conjugative plasmids [77]. A resident RM-carrying plasmid can exclude incoming plasmids with non-methylated restriction sites [78, 79]. In the case of co-infecting, incompatible plasmids, post-segregational killing will also introduce an advantage for the plasmid with the RM system [41, 80]. On the other hand, co-infecting compatible plasmids with RM systems may improve each others conjugation success, by modifying restriction sites that would otherwise be targeted upon entry into a recipient with an RM system.

CRISPR-Cas are also used by bacteria to defend against mobile genetic elements (MGEs). They typically consist of a ‘library’ of DNA fragments from past MGE infections (called ‘spacers’), and a system that cleaves any of those sequences once they are found in the cell [76]. CRISPR arrays, isolated cas genes, and entire CRISPR-Cas have been found on plasmids [75, 76]. CRISPR Type IV systems are almost exclusively found on plasmids, and interestingly, their spacers exhibit a strong bias towards other plasmids, specifically the transfer genes of conjugative plasmids [75]. Such systems can keep competing plasmids from establishing in the cell. Importantly, plasmid and chromosomal CRISPR-Cas can acquire immunity to plasmids they were previously (co-)infected with, thus shaping future infection dynamics.

Trans-acting exclusion systems can be implemented as follows:

- They lower the chance of successful plasmid transfer to recipients already carrying a plasmid (i.e. *k_i,j_* < 1).
- Post-segregational host killing due to plasmid-borne RM systems can be modelled similar to a TA system (see above).

## 4 Model application

In this section, we examine the influence of modelled co-infection processes on plasmid diversity. Our aim is to provide qualitative conceptual insights; the scale of our parameters is therefore arbitrary (Supplementary Table S1). We begin by considering two ecologically indistinguishable plasmid variants. This means that parameters values are identical for both variants (*β_A_* = *β_B_* = *β_AB_*, *k_A,B_* = *k_B,A_*, etc.; Supplementary Table S1). Further, by fulfilling a specific set of requirements (see Supplementary Text 3), we ensure that the model structure does not implicitly introduce an ecological difference between the variants (‘structural neutrality’) [36].

### 4.1 Influence of model parameters on co-infection

We begin by providing an intuition for the link between various model parameters and plasmid co-infection states by exploring the parameter space for plasmid conjugation (*β_i_*), infection susceptibility (*k_i,j_*), partial segregation loss (*m_i_*) and plasmid cost (*c_i_*, defined here as a decrease in replication rate due to plasmid carriage: *ρ_i_* = *ρ*_0_(1 – *c_i_*)). We randomly sample these parameters 6100 times (Supplementary Table S1) and classify the population output at steady state into the following outcomes: ‘no plasmid’ (*P*_0_), ‘high co-infection’ (*P_AB_*) or ‘low co-infection’ (*P_A_* and *P_B_*). The frequencies of each class over the whole data set show by far the highest prevalence of high co-infection (Figure 2A).

**Figure 2:**
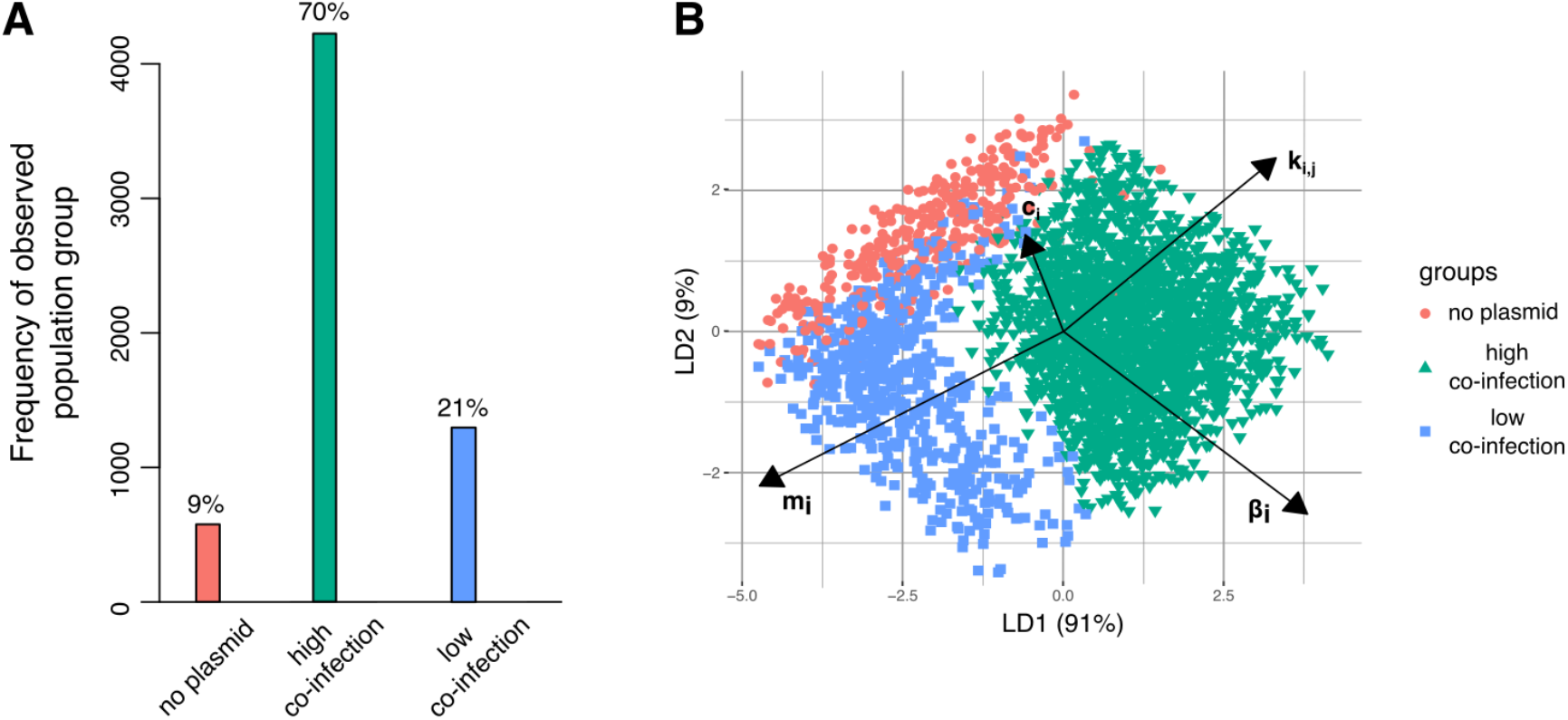
Parameter space exploration using linear discriminant analysis. **A**. Probability of each class over all simulation outcomes. Frequencies of each class at the end of 500 time steps - ‘no plasmid’ (red), ‘co-existence due to co-infection’ (green) or ‘co-existence without co-infection’ (blue) - are given for 6100 parameter sets randomly sampled over [0,0.5] for *m_i_* and [0,1] for *k_i,j_* = 2*k_i,AB_, β_i_, c_i_*. **B**. LDA using the 3 classes shown in A (same color scheme). Arrows show the magnitude and direction of the parameters varied (e.g. the shorter arrow of *c_i_* indicates lower significance of this parameter in class separation, whereas *m_i_* and *k_i,j_* (*k_i,AB_*) are most important in separating high from low co-infection areas).

Next, we identify the impact of each parameter on population dynamics using linear discriminant analysis (LDA). Briefly, LDA maximally separates the parameter regions, which tend to result in the different classes defined above [81]. We find that the most significant factors separating the two co-infection classes are susceptibility and partial segregation loss (as shown by the parameter arrows in Figure 2B), with increases in *k_i,j_* leading to more co-infections and increases in *m_i_* resulting in more single infections. The ‘no plasmid’ class is separated from the other two by low conjugation rates and high costs. While higher conjugation rates lead to more plasmid carriage in general, the direction of the arrow indicates that co-infections are relatively more increased. Notably, the magnitude of plasmid cost has the least influence on population outcome among these parameters, though this result may be sensitive to the overall parametrisation. On the whole, the co-infection parameters described here affect population outcomes in an intuitive and biologically meaningful manner.

### 4.2 Co-infection affects evolutionary outcomes through frequency-dependent selection

To explore the impact of co-infection on evolutionary outcomes, we again consider two ecologically indistinguishable plasmids variants. In a deterministic simulation, such indistinguishable competitors simply remain at their initial frequencies (Figure 3A). However, varying certain co-infection parameters (specifically, *ρ_AB_*, *γ_AB_*, *q_i_*, *β_AB_*, *g_i_* or the ratio between *k_i,j_* and *k_i,AB_*), while keeping all other parameter values identical for the two plasmid variants, changes plasmid dynamics by introducing frequency-dependent selection. This link between specific co-infection parameters and frequency-dependent selection is derived in Supplementary Text 3 and verified by simulation (Figure S2, Figure S3). The general insight (Figure 3A) is that frequency dependence arises from the impact of co-infection on the plasmid variants: when co-infection is beneficial for both co-residents, we observe negative frequency-dependent selection (NFDS); when it is detrimental to both variants, we observe positive frequency-dependent selection (PFDS).

**Figure 3:**
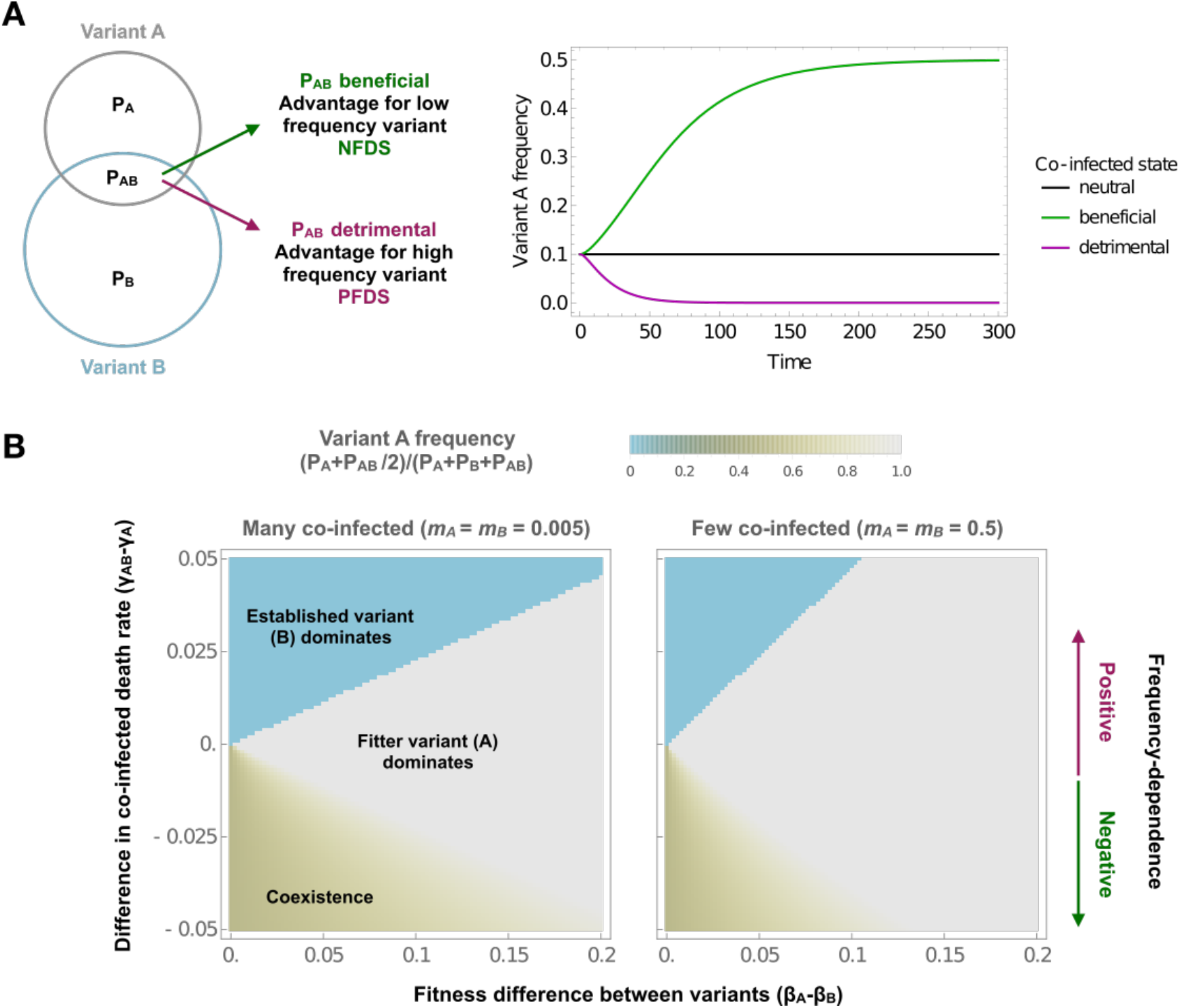
Co-infection affects evolutionary outcomes through frequency-dependent selection. **A**. The effect of the co-infected state on the outcome of competition between two plasmid variants with identical properties. When the co-infected state is neither beneficial nor detrimental, there is no frequency-dependent selection and the plasmid variants remain at their initial frequencies. A co-infection related advantage for both variants introduces negative frequency-dependent selection, which equalises variant frequencies and leads to co-existence. A co-infection related disadvantage introduces positive frequency-dependent selection, which leads to the exclusion of the variant with a lower initial frequency. **B**. The effect of frequencydependent selection on evolutionary outcomes in presence of fitness differences. The figures show the equilibrium frequency of a variant with a fitness advantage but with low initial frequency (*P_A_* = 0.001 and *P_B_* = 1 at *t* = 0). The color indicates the equilibrium frequency of variant A (here defined as *P_A_* + *P_AB_*/2 at *t* = 300000). The x-axis captures the extent of the fitness difference, here implemented as a difference in conjugation rate (*β_i_*). The y-axis captures the strength and direction of the frequency-dependent selection, here implemented by varying the death rate (*γ_i_*) of the co-infected cells. For both plots, standard parameter values are: *ρ*_0_ = 1, *ρ_A_* = *ρ_B_* = *ρ_AB_* = 0.9, *γ_i_* = 0.1, *β_A_* = *β_B_* = 0.2, *β_AB_* =0, *m_i_* = 1/3, *q_i_* = 1/2, *s_i_* = 1/1000 *k_A,B_* = *k_B,A_* = 1/2, *k_A,AB_* = *k_B,AB_* = 1/4

This frequency-dependence arises because the frequency of a plasmid variant determines the relative contribution of the co-infected state to its overall reproductive success, which depends on both *P_A_* (*P_B_*) and *P_AB_*. If variant A is rarer than variant B (*P_A_* < *P_B_*), the co-infected state makes up a larger proportion of the overall density of plasmid A (*P_AB_*/(*P_AB_* + *P_A_*) > *P_AB_*/(*P_AB_* + *P_B_*)). Therefore, if the co-infected state is beneficial for both plasmids, rare variants have an advantage, which will equalise variant frequencies. Conversely, if the co-infected state is detrimental, rare variants have a disadvantage, allowing the variant with a higher initial frequency to exclude the other. Intuitively, the co-infected state is beneficial when co-infected cells have a higher net growth rate; an overall higher conjugation rate; a lower probability of complete segregation loss; or are less susceptible to further infection with other plasmids (Supplementary Text 3).

Next, we explore the effect of introducing a fitness difference between the plasmids (Figure 3B). As expected, both NFDS and PFDS can lead to persistence of the lower fitness variant: NFDS by allowing co-existence of the two competitors, and PFDS by preventing the higher fitness variant from invading a population in which the lower fitness variant is already established. In both cases, whether the lower fitness variant is maintained depends on the strength of the frequency-dependent selection relative to the fitness difference. The frequency-dependent effect is stronger when co-infection is common. Thus, parameters which do not themselves introduce frequency-dependent selection but affect the frequency of the co-infected state (e.g. *m_i_* and *k_i,j_*), can influence evolutionary outcomes by modulating the strength of frequency-dependent effects.

Finally, we consider the impact of asymmetric co-infection related effects. Thus far, we analysed effects which are equally beneficial or detrimental for both co-infecting variants: either because they impact properties of the host cell (e.g. *rho_AB_*), or because we have assumed within-host interactions to be symmetric (e.g. *q_A_* = *q_B_*, *m_A_* = *m_B_*,...). However, within-host interactions can also be asymmetric (see Section 3): for example, between incompatible plasmids, an advantage in replication and/or partitioning would translate to a difference in partial segregation loss (*m_i_* < *m_j_*) and conjugation from co-infected cells (*q_i_* > *q_j_*) through changes in within-cell variant frequencies. Such asymmetric effects give one of the variants a competitive advantage (Figure S4), but do not, in themselves, introduce frequency-dependent effects (Supplementary Text 1.4).

## 5 Discussion

This work provides an overview of the biological processes relevant in plasmid co-infection (Section 3) and discusses how they can be captured appropriately in a modeling framework (Section 2). We demonstrate how this general framework can be applied to understand how co-infection parameters shape plasmid variant selection and diversity.

Depending on the underlying processes, the co-infected state can be either beneficial or detrimental for the plasmid variants. Benefits arise for example from ‘collaborative’ (i.e. higher overall) conjugation from co-infected cells; positive epistasis in host fitness (reduced plasmid cost or higher plasmid benefit); or distinct cis-acting exclusion systems (protecting the cell from further infection with either variant). We would therefore expect negative frequency-dependent selection to maintain diversity in these traits. Conversely, with negative epistasis or addiction systems co-infection would be detrimental, making displacement of established variants difficult due to positive frequency-dependent selection. Finally, replication or partitioning incompatibility does not in itself lead to frequency-dependent selection, but does modulate its strength by decreasing the density of co-infected cells.

These co-infection related effects also have implications on the evolutionary trajectories of bacterial populations more broadly. Co-infection influences the rate at which bacterial populations acquire new genes through plasmid transfer: the entry of plasmids from other bacterial cells or species is influenced by the presence of a resident plasmid [7, 17]. By promoting the introduction of new variants, negative frequency dependence can act to increase the acquisition of plasmids from other bacterial populations/species. Conversely, positive frequency dependence can act as a barrier to new plasmids entering the population, thus slowing this acquisition. Secondly, co-infection governs the extent of plasmid gene sharing. When present in the same cell, plasmids can exchange genetic material through e.g. recombination [64, 82]. Frequency-dependent effects would also be expected to influence the mobility of genes between plasmids (or plasmid and chromosome [83]). For example, if the presence of the same cargo gene on (compatible) co-resident plasmids gives rise to negative epistasis between the plasmids (due to negative gene dosage effects), the resulting PFDS would constrain gene mobility: the disadvantage associated with low frequency variants would prevent plasmids that have newly acquired the cargo gene from increasing in frequency.

Our results are closely linked to previous theoretical work on epidemiological models of co-infection [36], which has highlighted how model structure can include coexistence-promoting mechanisms. Specifically, the motivating concern of this previous work was that models of co-infection typically implicitly and unintentionally assumed that a host carrying one strain would be susceptible to co-infection with another strain, but protected from re-infection with itself: co-infection was possible, but displacement was neglected. This is akin to assuming cis-acting exclusion. In models of plasmid co-infection, this specific concern is, to some extent, less acute cis-acting exclusion systems are thought to be widespread among conjugative plasmids [7]. If these systems are indeed as effective *in vivo* as *in vitro* data suggest, co-infection only occurs between plasmids of different exclusion groups and co-infected cells are therefore indeed not susceptible to displacement. Furthermore, when considering variants of the same backbone with and without a particular cargo gene, it is appropriate to exclude co-infection [11, 83]. On the other hand, our results highlight that frequency-dependent effects also arise from other model features. Many of these effects are linked to copy number, making evolutionary outcomes heavily dependent on how co-infecting plasmids influence each others’ copy numbers. It is thus important to be explicit about the traits of the modelled variants and aware that results may not generalise for different assumptions about plasmid backbones.

A key feature of the framework discussed here is that cells are tracked in terms of the plasmid variants they carry, without explicitly incorporating plasmid copy number: each cell type (*P*_0_, *P_A_*, *P_B_*, *P_AB_*) is represented in terms of the average cell, and heterogeneity within cell types is ignored. This is a standard approximation in compartmental models, but warrants additional discussion in the context of co-infection. Firstly, this approximation can make the link between model and biological processes less intuitive and complicates parametrisation, as processes which change within-cell plasmid frequencies have to be represented in terms of average plasmid loss. Secondly, by representing the co-infected state as a single variable, the average frequency of plasmid variants within co-infected cells is implicitly specified. This highlights the importance of carefully considering how certain parameters values depend on relative plasmid frequencies (e.g. *k, m, q*), particularly when modelling plasmids where one variant has a within-cell competitive advantage and thus the variant frequencies within-co-infected cells are not equal. Overall, the contexts in which explicit models of plasmid copy number are not satisfactorily approximated by average copy numbers warrants further exploration (Supplementary Text 2.2).

To truly understand the eco-evolutionary implications of co-infection, more empirical research is needed on its natural occurrence and distribution. This includes population-level studies investigating the prevalence of plasmid co-infection across bacterial phyla, as well as its correlation with incompatibility group, plasmid size, and copy number. Further, while studied in detail at the mechanistic level, little is known about the natural diversity and phenotypic effects of various exclusion and TA systems. Carefully designed bioinformatics studies could address some of these questions. However, sequencing databases are currently not representative of natural microbial diversity, and the meta-data to account for phylogenetic, geospatial, or phenotypic biases is often lacking [84]. Additionally, plasmids may not be represented accurately in the deposited genomes [85, 86], complicating conclusions on overall plasmid co-infection.

A combination of empirical and theoretical approaches is necessary to iteratively refine our understanding of plasmid diversity: on the one hand, using empirical data to inform model parameter values and processes, and on the other, evaluating the results of simulations against natural observations. In particular, combining insights into the mechanistic effects of specific traits from experimental studies and data on the distribution of these traits in natural plasmid populations is a crucial step, and modelling can provide an important tool in bridging these two levels of observation. Through careful consideration of the biological processes and potential modelling pitfalls relating to plasmid co-infection, we have developed a modelling framework which can serve as a basis for such future work.

## 1 Supplementary Methods and Results

### 1.1 Parameter table for simulations

**Table S1:**
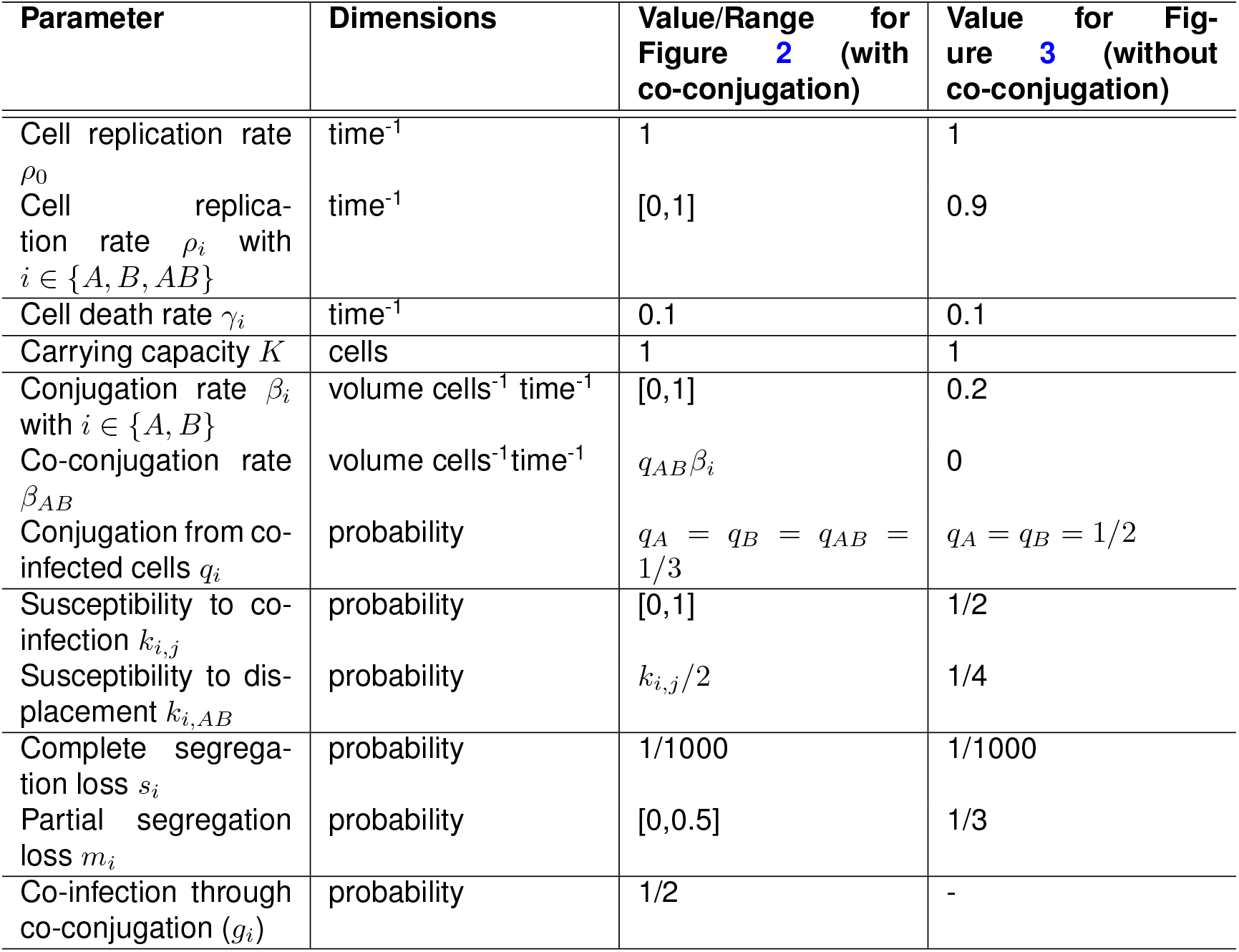
Parameter dimensions and values or ranges used in simulations. For these parameter values, the model is structurally neutral - i.e. the variants are ecologically indistinguishable (see Supplementary Text 3). Unless otherwise indicated, the values for *A, B* and *AB* are the same in all simulations. The two columns represent parameters used in simulations with and without co-conjugation (i.e. simultaneous transfer of both plasmids from co-infected cells). Most simulations relating to frequency-dependent selection (e.g. Figures 3 and S2) use the parameter set without co-conjugation because the relationship with previous work on structural neutrality is clearer when *β_AB_* = 0. Frequency-dependent selection with co-conjugation is explored in Figure S3. Co-infection through co-conjugation (*g_i_*) is not relevant when there is no co-conjugation, which is why the value is in the third column.

| Parameter | Dimensions | Value/Range for Figure 2 (with co-conjugation) | Value for Figure 3 (without co-conjugation) |
| --- | --- | --- | --- |
| Cell replication rate $\rho_0$ | time<sup>-1</sup> | 1 | 1 |
| Cell replication rate $\rho_i$ with $i \in \{A, B, AB\}$ | time<sup>-1</sup> | [0,1] | 0.9 |
| Cell death rate $\gamma_i$ | time<sup>-1</sup> | 0.1 | 0.1 |
| Carrying capacity $K$ | cells | 1 | 1 |
| Conjugation rate $\beta_i$ with $i \in \{A, B\}$ | volume cells<sup>-1</sup> time<sup>-1</sup> | [0,1] | 0.2 |
| Co-conjugation rate $\beta_{AB}$ | volume cells<sup>-1</sup> time<sup>-1</sup> | $q_{AB}\beta_i$ | 0 |
| Conjugation from co-infected cells $q_i$ | probability | $q_A = q_B = q_{AB} = 1/3$ | $q_A = q_B = 1/2$ |
| Susceptibility to co-infection $k_{i,j}$ | probability | [0,1] | 1/2 |
| Susceptibility to displacement $k_{i,AB}$ | probability | $k_{i,j}/2$ | 1/4 |
| Complete segregation loss $s_i$ | probability | 1/1000 | 1/1000 |
| Partial segregation loss $m_i$ | probability | [0,0.5] | 1/3 |
| Co-infection through co-conjugation ($g_i$) | probability | 1/2 | - |

### 1.2 Classification using linear discriminant analysis (LDA)

We explored the influence of various co-infection parameters by random parameter sampling and subsequent separation through linear discriminant analysis (LDA) for the population dynamics of two identical plasmids, i.e. where all infection and co-infection parameters between the plasmids are the same. The only difference we assume between the plasmids is the starting frequency, which is higher for plasmid variant *B*. We use two different sets of initial conditions: either starting from an almost entirely susceptible population (94% *P*_0_, 1% *P_A_*, 5% *P_B_*) or starting from an almost entirely plasmid-carrying population (98% *P_B_*, 1% *P_A_*, 1% *P*_0_). (The results of the former are shown in the main text (Figure 2) and for the latter in Figure S1.)

First, we randomly sampled the parameter space (6100 samples) for the following parameters: plasmid transmission (*β_i_*), susceptibility to co-infection or displacement (*k_i,j_, k_i,AB_*), partial plasmid loss (*m_i_*) and plasmid cost (i.e. decreases in replication rate *ρ_i_*) using linear sampling between 0 and 1 or 0 and 0.5 (Table S1). We recorded the frequency of each subpopulation after 500 time steps (we also used 1000 time steps to ensure that our simulations had reached steady state and saw no difference in the outcome) and used these frequencies for classification of the outcome into 4 classes:

- ’no plasmid’: *P*_0_/(*P*_0_ + *P_A_* + *P_B_* + *P_AB_*) > 90%
- ’one plasmid’: *P_A_*/(*P_A_* + *P_B_* + *P_AB_*) > 50% or *P_B_*(*P_A_* + *P_B_* + *P_AB_*) > 50%, and all other populations < 20%
- high co-infection’: *P_AB_*(*P_A_* + *P_B_* + *P_AB_*) > 50%, and all other populations < 20%
- ’low co-infection’: *P_A_* and *P_B_* each > 25% and together > 50%, and all other populations < 20%

This classification was used to identify the effect of each parameter on the population outcome by applying linear discriminant analysis (LDA), which is a supervised method of dimensionality reduction [1]. Specifically, LDA projects the simulation results on a 2D space so that the centroids of the individual classes are maximally separated and the within-class scattering of points is minimized. The magnitude and direction of the parameter arrows in this 2D-space show their significance in separating specific classes.

The ‘one plasmid’ class is observed at a very low probability in our data set (Figure S1A): we assume equal fitness for both plasmids and this class therefore reflects the influence of initial conditions (i.e. occurs when the low initial frequency plasmid remains at low frequency). Accordingly, there is significant overlap between the ‘one plasmid’ and ‘low co-infection’ class (Figure S1B). This makes sense as the parameters explored here are mostly modifying co-infection behavior and will not be instrumental in separating variations in singly infected states (i.e. *P_A_* and *P_B_* frequencies). Hence, we are not showing the ‘one plasmid’ class for the LDA in Figure 2, but the results are highly similar to the ones shown in Figure S1A, B.)

**Figure S1:**
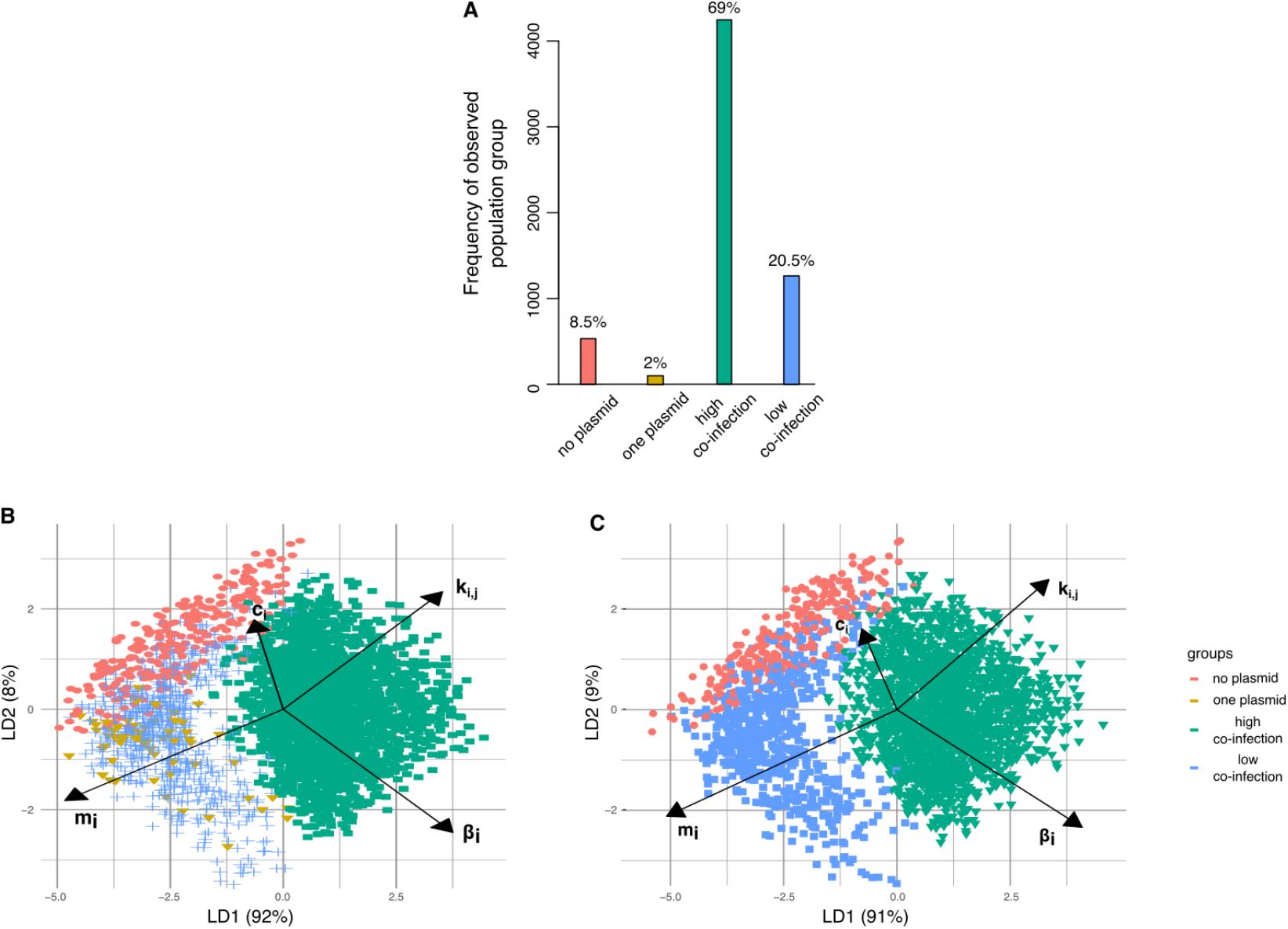
LDA performed on different initial conditions. **A**. Probability of each class over all simulation outcomes. Frequencies of each class - ‘no plasmid’ (red), ‘one plasmid’ (yellow), ‘high co-infection’ (green) or ‘low co-infection’ (blue) - are given for 6100 randomly sampled parameter sets (*m_i_,k_i,j_* = 2*k_i,AB_,β_i_,c_i_*) at the end of 500 time steps. **B**. LDA using all 4 classes shown in A (using the same color scheme). Arrows show the magnitude and direction of the parameters varied. The survival of only one plasmid cannot be well separated from the other classes with these parameters. **C**. LDA without the ‘one plasmid’ class. Parameters *m_i_* and *k_i,j_* (*k_i,AB_*) are most important in defining separability of plasmid co-existence due to co-infection or singly infected cells.

### 1.3 Parameter values and frequency-dependent selection

Figures S2 and S3 show how deviation from the parameter values in Table S1 introduces frequency-dependent selection, for a certain subset of parameters. Specifically, the variants are under NFDS (i.e. co-infection is beneficial) when:

- co-infected cells have a higher replication rate than the singly infected state (*ρ_AB_* > *ρ_A_*, or, equivalently, *ρ_AB_* > *ρ_B_*);
- co-infected cells have a a lower death rate (*γ_AB_* < *γ_A_*);
- co-infected cells have a lower probability of plasmid loss (*s_AB_* < *s_A_*);
- the overall rate of plasmid transmission from co-infected cells is higher 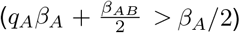;
- co-conjugation to a singly infected cell has a higher probability of resulting in co-infection than expected 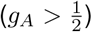;
- if a plasmid in *P_AB_* is less susceptible to further infection by the competing variant 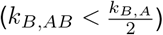.

Reversing these inequalities makes the co-infected state detrimental and leads to PFDS. Changes in other parameters (*β_i_, m_i_, k_i,j_*) do not introduce frequency-dependent effects (see Section 3 for mathematical derivation).

**Figure S2:**
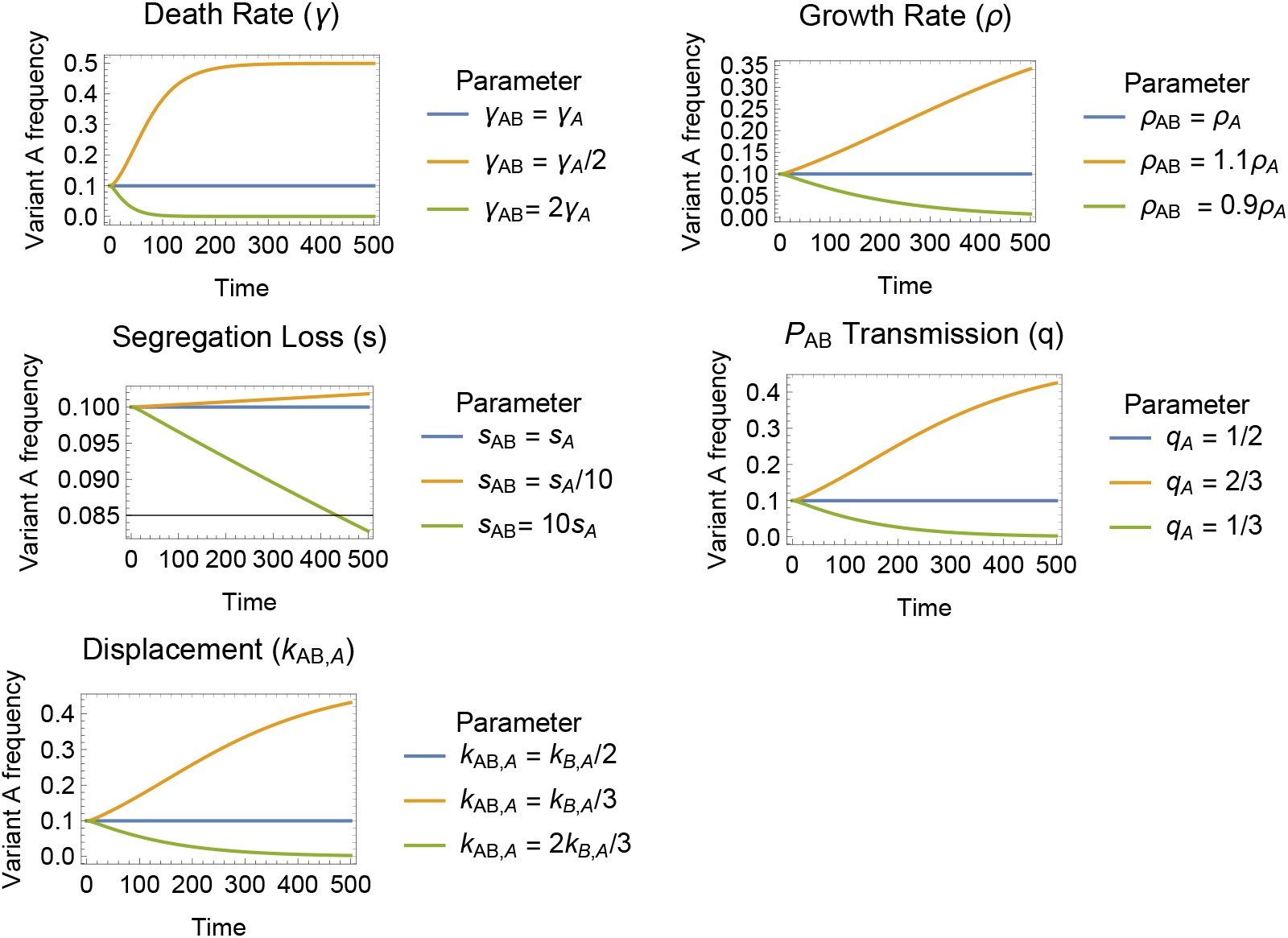
The impact of model parameters on frequency-dependent effects in a model without co-conjugation (*β_AB_* = 0) Each plot shows the effect of varying a single parameter. For all plots, the properties of both plasmid variants are always identical. Standard parameter values are as follows: *ρ*_0_ = 1, *ρ_A_* = *ρ_B_* = *ρ_AB_* = 0.9, *γ_i_* = 0.1, *β_A_* = *β_B_* = 0.2, *β_AB_* = 0, *m_i_* = 1/3, *q_i_* = 1/2, *s_i_* = 1/1000 *k_A,B_* = *k_B,A_* = 1/2, *k_A,AB_* = *k_B,AB_* = 1/4.

### 1.4 Asymmetric co-infection effects

Figure S4 shows the impact that asymmetric co-infection effects (specifically, partial segregation loss, *m_i_*, and horizontal transmissibility from co-infected cells, *q_i_*) have on model outcomes. These asymmetries do not in themselves introduce frequency-dependent selection, although with horizontal transmissibility (*q_i_*), frequency-dependent effects arise from changes in the overall conjugation rate from co-infected cells.

**Figure S3:**
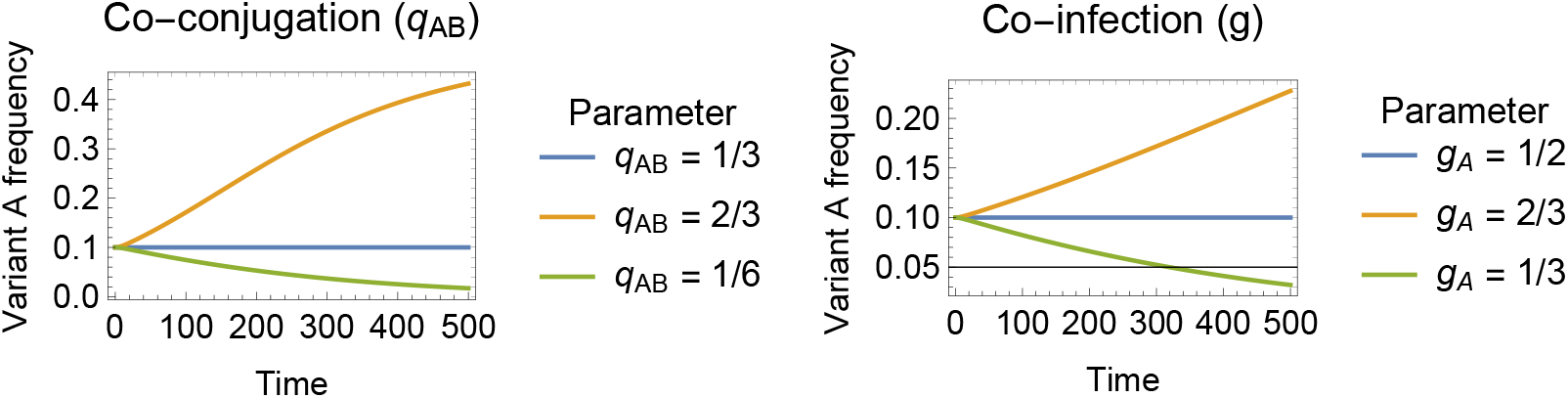
The impact of parameters relating to co-conjugation on frequency-dependent effects. For this plot, we have allowed co-conjugation (*β_AB_* = *q_AB_β_A_* = *q_AB_β_B_*) and reduced the q parameters (*q_A_* = *q_B_* = *q_AB_* = 1/3). Each panel explores the effect of parameters relating to co-conjugation. The properties of both plasmid variants are always identical. Other default parameter values are: *ρ*_0_ = 1, *ρ_A_* = *ρ_B_* = *ρ_AB_* = 0.9, *γ* = 0.1, *β_A_* = *β_B_*, *m_i_* = 1/3, *s_i_* = 1/1000 *k_A,B_* = *k_B,A_* = 1/2, *k_A,AB_* = *k_B,AB_* = 1/4, *g* = 1/2.

**Figure S4:**
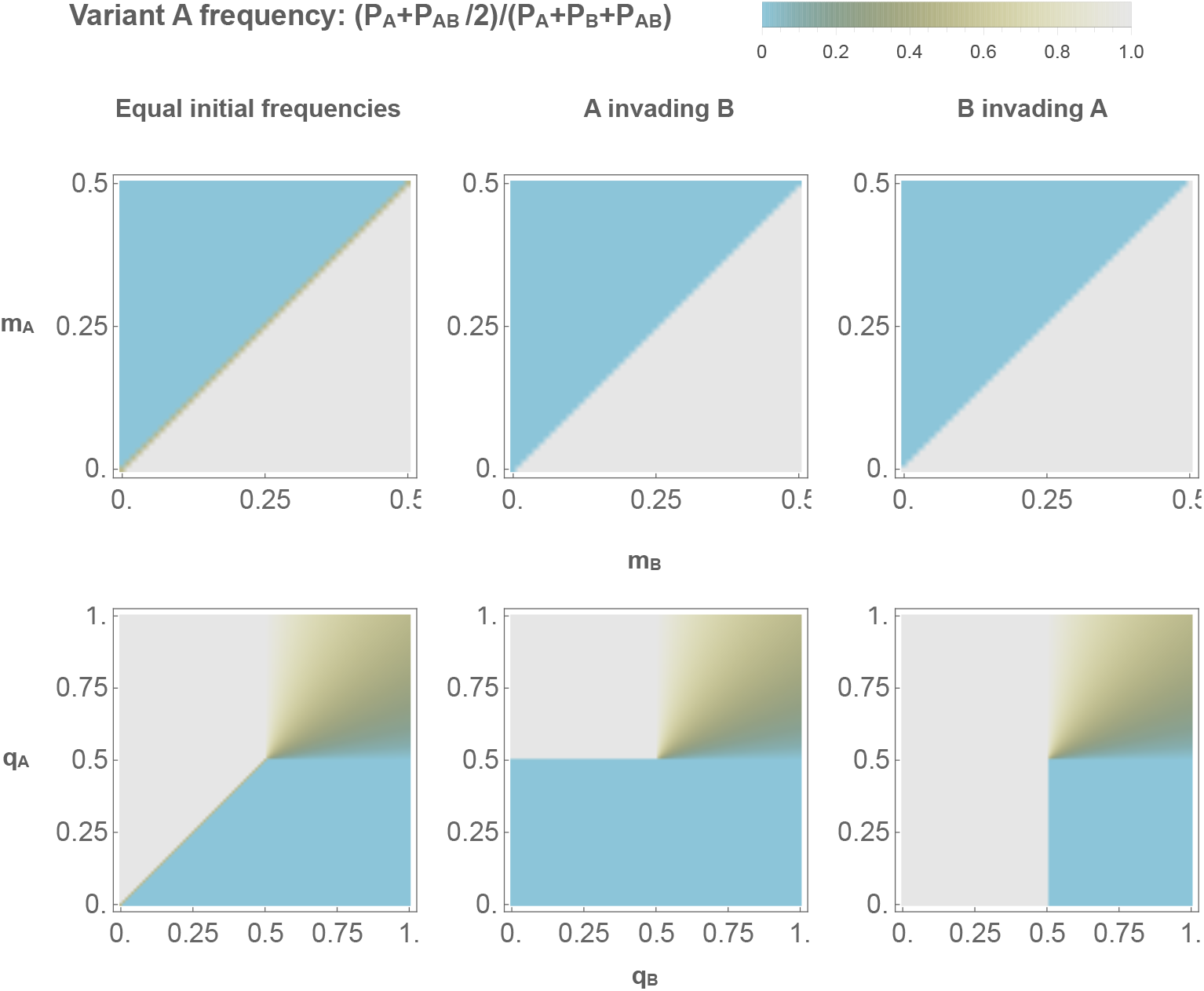
The effect of asymmetric co-infection interactions on frequency-dependent selection. The colour in each plot indicates the equilibrium frequency of variant *A*. The columns indicate different initial plasmid variant frequencies. Top row: the effect of asymmetric probability of vertical transmission for co-infecting plasmids (*m_i_* indicates the probability that plasmid variant *i* is lost during replication). No frequency-dependent effects are observed. Bottom row: the effect of an asymmetric probability of horizontal transmission for co-infecting plasmids (*q_i_* indicates the probability that variant *i* is transmitted during conjugation by coinfected cells). Frequency-dependent effects do not arise when the overall transmissibility of the co-infected cell is the same as of singly infected cells: when *q_A_* + *q_B_* = 1, (i.e. the main diagonal of each plot), the fitter variant (i.e. with higher *q_i_*) always out-competes the other, regardless of initial conditions. For all plots, standard parameter values are as follows: *ρ*_0_ = 1, *P_A_* = *P_B_* = *P_AB_* = 0.9, *γ_i_* = 0.1, *β_A_* = *β_B_* = 0.2, *β_AB_* = 0, *m_i_* = 1/3, *q_i_* = 1/2, *s_i_* = 1/1000 *k_A,B_* = *k_B,A_* = 1/2, *k_A,AB_* = *k_B,AB_* = 1/4

## 2 Considerations around model structure

Here, we develop some ideas relating to model structure in more depth.

### 2.1 The nature of co-infection

In drawing parallels between infectious disease and plasmid dynamics, it is worth making a distinction between two fundamentally different forms of co-infection (Figure S5A). In the first form of co-infection, hosts contain multiple ‘patches’: in singly infected hosts, a single patch is occupied, and co-infection occurs through additional patches becoming occupied. In this form of co-infection, a host can be multiply infected with the same variant, and this state is ecologically different from single infection with that variant. This type of co-infection can be appropriate for epidemiological models – e.g. when co-infection represents infection of multiple body sites – but it is difficult to see a biological correspondence of different within-cell patches in plasmid infection.

In the second form of co-infection, which is appropriate for modelling plasmid co-infection, hosts consist of a single patch, which can be occupied by one or more variants. In this single patch form of co-infection, multiple infection with the same variant is indistinguishable from single infection with that variant. Cells which are multiply infected with different variants may have different properties from singly infected cells, such as higher overall within-host copy number, for example. The key distinction is that this difference arises from the properties of co-infecting variants–e.g. through dependence on different within-host resources–and is not an inherent property of the host cells. This distinction is relevant when relating our results to the concept of structural neutrality in epidemiological models (SI Section 3). [2].

### 2.2 Implicit modelling of plasmid copy number

The type of model we discuss does not explicitly track plasmid copy number (Figure S5B): for example, the entire *P_A_* population is approximated by a single cell type representing the average copy number, ignoring stochastic copy number variation as well as copy number dynamics during the cell cycle and following initial infection. In general, this is likely to be a reasonable approximation as copy number adjustments are generally very fast, with plasmid number doubling times of 5-10 minutes [3]. In models of plasmid co-infection, this approximation also means that we do not explicitly model within-cell variant frequencies. The co-infected state represents the average co-infected state, i.e. average within-cell variant frequencies. If the average within-cell variant frequencies are expected to be equal (i.e. 50/50 on average), copy number dependent parameters (e.g. *k_i,AB_, m_i_, q_i_*) would also be equal for both variants (in absence of other copy number independent effects that could introduce a difference in these parameters). Unequal average within-cell frequencies can be expressed by adjusting copy number dependent parameters to reflect the greater copy number of one variant. Here, it is important to be explicit about the assumed relationship between copy number and parameter value. Overall, although the implicit modelling of copy number does not necessarily affect evolutionary outcomes, it makes the relationship between biological processes and model parameters less intuitive.

### 2.3 Cell population growth and resource competition

We have modelled net cell population growth as consisting of both a density-dependent and density-independent component. It is worth highlighting that growth can also be modelled without inclusion of a density-independent element, i.e. with a single term relating net population growth to the carrying capacity *K*. Both versions of the model lead to a bounded total population size. The essential difference is that, without the density-independent term, cells neither replicate nor die once carrying capacity is reached. In this version of the model, therefore, the effect of plasmid carriage on host cell fitness ceases to matter once carrying capacity is reached. In terms of evolutionary outcomes, the two types of model will place a different emphasis on horizontal and vertical effects. In cases where the fitness difference between plasmid variants arises solely through effects on the host cell fitness (e.g. competition between variant with and without a specific cargo gene, such as antibiotic resistance), this difference is substantial: a model without a density-independent growth term will generally allow coexistence of the variants at equilibrium, if carrying capacity is reached before one of the variants has out-competed the other.

**Figure S5:**
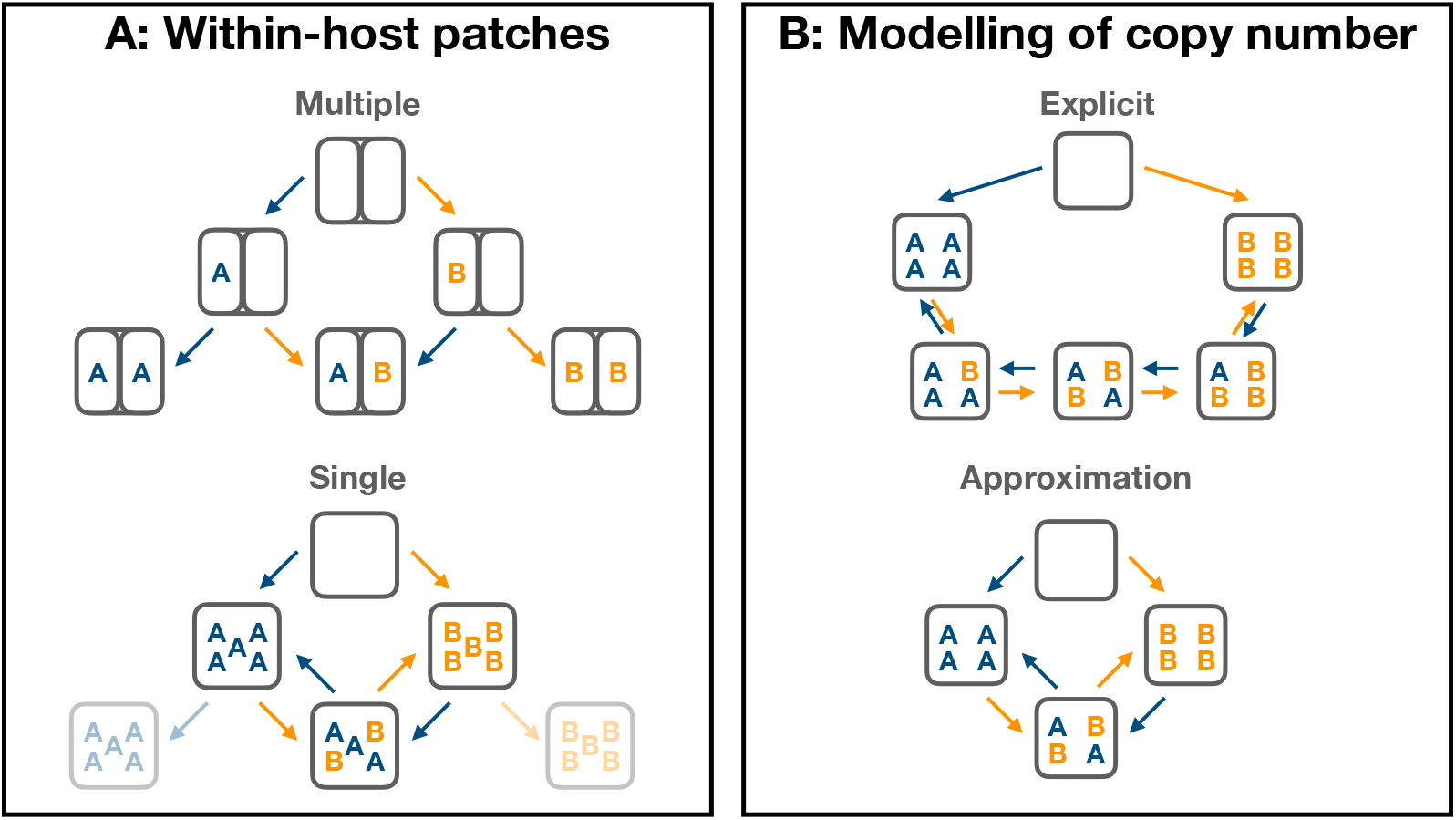
Illustration of two concepts central to modelling plasmid co-infection. **A** Coinfection with and without within-host patches. In the top diagram, hosts consist of multiple patches, and co-infection occurs when more than one patch is occupied. In the bottom diagram, hosts consist of a single patch and co-infection occurs when this patch is shared by multiple variants. An important difference between the two types of co-infection is that co-infection with the same variant is not biologically meaningful in the single patch model (illustrated as transparent states in the lower panel). **B** Explicit and implicit modelling of copy number. The top diagram illustrates what a model explicitly representing within-host variant frequencies might look like. The bottom diagram shows how this explicit model is approximated.

## 3 Co-infection and frequency-dependent selection

Here, we discuss the relationship between co-infection and frequency-dependent selection in more depth and consider this result in relation to Lipsitch et al. [2], which motivated much of our approach. Specifically, we discuss the concept of structural neutrality in greater depth; use this approach to derive which parameters lead to frequency-dependent effects; and expand on the interpretation of these effects in terms of beneficial and detrimental effects of co-infection.

### 3.1 Structural neutrality

Work on epidemiological models of co-infection has shown that seemingly innocuous modelling choices can introduce unintended ecological differences between strains, which can act to promote strain diversity (‘co-existence for free’) [2]. To ensure that model outcomes reflect the intended ecological properties of strains, rather than unintended differences arising by model structure, Lipsitch et al. introduced the concept of a structurally neutral null model. Such null models satisfy criteria (discussed below) which ensure they do not contain unintended ecological interactions and can thus serve as a starting point for introducing intended differences. For example, consider a model of competition between an antibiotic resistant and an antibiotic susceptible strain: the intended differences between the strains would arise from the effect of antibiotics and a potential fitness cost of resistance. Thus, in absence of antibiotic pressure and with no resistance-associated fitness cost, the two strains are ecologically indistinguishable and the model structure should reflect this.

Lipsitch et al. define structural neutrality in a model of indistinguishable strains (i.e. strains, or in our case plasmids, differing only in a neutral marker) in terms of two criteria. Firstly, ‘ecological neutrality’: if the strains are indeed indistinguishable, they should have identical ecological interactions, i.e. there should be nothing distinguishing the interactions that strain A has with itself, from the interactions that strain A has with strain B. For this criterion to hold, it should be possible to write the dynamics of the number of strains a host is infected with (referred to as ‘ecological variables’ in Lipsitch et al.), without reference to particular strain identities. For example, in our case, the ecological variables would be defined by the number of plasmids variants in the host cell, i.e. *P*_0_, *P*_1_ (where *P*_1_ = *P_A_* + *P_B_*), and *P*_2_ (where *P*_2_ = *P_AB_*). For the ecological neutrality criterion hold, it should be possible to write the equations for these ecological variables without making reference to *P_A_* and *P_B_*. The intuition behind this criterion is that, because *A* and *B* are indistinguishable, the dynamics of *P*_0_, *P*_1_ and *P*_2_ should not depend on how *P_A_* and *P_B_* make up the *P*_1_ class.

The second criterion is ‘population genetic neutrality’: there should be no single equilibrium strain frequency, instead, any arbitrary equilibrium strain frequency should be reachable by choosing the right initial conditions. The intuition for this criterion is that, with two indistinguishable strains, there should be no mechanism that acts to equilibrate strain frequencies. Lipsitch et al. show that models which can be written in a specific form (‘ancestor-tracing’, see Lipsitch et al.) that fulfills the ecological neutrality criterion also fulfill the population genetic one.

### 3.2 Relationship between our results and structural neutrality

Our results are closely related to the concept of structural neutrality: for parameter values at which the co-infected state is neither beneficial nor detrimental for the plasmid variants, the model is neutral. Figure 3A and Supplementary Figures S2 and S3 illustrate population genetic neutrality: this is equivalent to the absence of positive or negative frequency-dependent selection.

As written, the model does not directly fulfill the ecological neutrality criterion: the dynamics of *P*_1_ are not independent of variant identities (i.e. the equation for 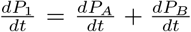 cannot be written just in terms of the ecological variables *P*_0_, *P*_1_ and *P*_2_, but remains dependent on *P_A_* and *P_B_*). However, it is possible to re-formulate the model in a mathematically equivalent form, at least when *β_AB_* = 0, which does fulfill the ecological neutrality criterion (as demonstrated in Lipsitch et al. [2]). To achieve this, we need to introduce two additional states, where a host is dually infected with the same variant (*P_AA_* and *P_BB_*). As discussed in Supplementary Text 2.1, if the host consists of a single patch, these states are not biologically meaningful. However, we can introduce them as a mathematical convenience to produce a model which does fulfill the ecological neutrality criterion (see Lipsitch et al. for an ecological interpretation of this alternative formulation). Switching to *N* to denote cell densities in order to facilitates comparison between the two formulations, we consider a model of the form:

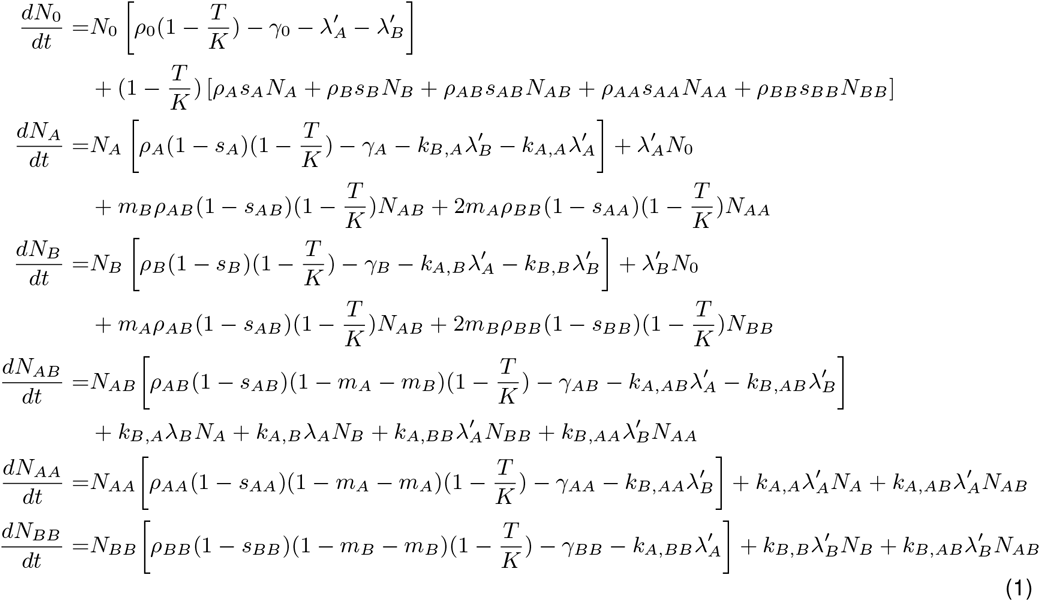

Here, 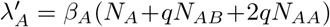 and 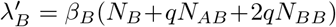. With *P_A_* = *N_A_* + *N_AA_* and *P_B_* = *N_A_* + *N_BB_*, this model is identical to the main text model (Equations 1, assuming *β_AB_* = 0) when:

- *q* = 1/2
- *k_A,B_* = *k_A,BB_* = 2*k_A,AB_* and *k_B,A_* = *k_B,AA_* = 2*k_B,AB_*
- *ρ_A_* = *ρ_AA_* and *ρ_B_* = *ρ_BB_*
- *γ_A_* = *γ_AA_* and *γ_B_* = *γ_BB_*
- *s_A_* = *s_AA_* and *s_B_* = *s_BB_*

Most of these criteria are straightforwardly interpretable: the two model formulations are equivalent when the *N_i,i_* state is identical to the *N_i_* state. The intuitive explanation for the criteria relating to the *k* parameters is discussed below.

In addition, this model fulfills the ecological neutrality criterion (i.e. the dynamics of the ecological variables *N*_0_, *N*_1_ = *N_A_* + *N_B_* and *N*_2_ = *N_AA_* + *N_BB_* + *N_AB_* can be written without reference to the specific variant identities) when all parameters are identical for plasmid *A* and *B* and:

- *ρ_AB_* = *ρ_AA_* = *ρ_BB_*
- *γ_AB_* = *γ_AA_* = *γ_BB_*
- *s_AB_* = *s_AA_* = *s_BB_*

These criteria are straightforwardly interpretable as the NAB state being equivalent to the *N_AA_* and *N_BB_* (and hence *N_A_* and *N_B_*) states. Thus, when both sets of criteria hold, the co-infected state is equivalent to the singly infected state, and the model is structurally neutral. Thus, this reasoning recovers the results from Supplementary Figure S2 and the interpretation of neutrality arising from the co-infected state being neither beneficial nor detrimental. Note that these criteria do not include any specific constraints on *k_A,B_* and *k_B,A_* (other than in relation to *k_A,AB_* and *k_B,AB_*) or *m_A_* and *m_B_*.

In a model which includes co-conjugation (*β_AB_* > 0), the re-formulation to demonstrate structural neutrality is more cumbersome and beyond the scope of this paper. However, intuitively (and as verified by simulation in Supplementary Figure S3) the co-infected state is equivalent to the singly infected state (and the model is neutral) when:

- the overall infectiousness of the two state is the same: 2*qβ_A_* + *β_AB_* = *β_A_* (and similarly for *B*)
- co-conjugation leads to co-infection half as often as single conjugation (see below for further explanation): *g_A_* = *g_B_* = 1/2.

Finally, a small technical note: Lipsitch et al. consider ‘closed’ transmission models specifically, where all infected individuals were either infected at time 0 or result from horizontal transmission. Technically, our model of plasmid transmission, and models of plasmid dynamics more generally, do not fall into this category because these models also allow infected cells to arise through replication of infected cells (i.e. vertical transmission). However, the results in Lipsitch et al. are nevertheless applicable to these models as well: the reasoning in Lipsitch et al. requires models to be closed to ensure that all strains present in the system had an ‘ancestor’ present at time 0 (as opposed to coming into the system through migration, for example). This ancestry criterion holds for models allowing for vertical transmission, as each infected cell remains traceable to an ancestor infection.

### 3.3 Interpreting the impact of co-infection

The section above develops a mathematical explanation for why changes in specific parameters lead to frequency-dependent effects. In this section, we elaborate on the biological explanation, i.e. which biological processes make the co-infected state beneficial or detrimental to the co-infecting plasmids. For most parameters (Supplementary Text 1.3), the effect of the co-infected state is intuitive: for example, the co-infected state is clearly beneficial for co-infecting plasmids if co-infected cells have a lower death rate; a lower probability of overall plasmid loss; or higher overall conjugation rate than singly infected cells.

The results regarding susceptibility to co-infection and displacement (i.e. co-infection is beneficial when 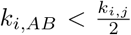 and 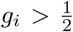) are perhaps a little less intuitive. For the susceptibility to displacement (*k_i,AB_*) it is helpful to consider a plasmid of copy number two. When establishing co-infection (*P_A_* → *P_AB_*), the incoming plasmid *B* can replace either of the copies of plasmid *A*, whereas displacement (*P_AB_* → *P_B_*) only occurs if the incoming plasmid *B* replaces the *A* variant. The reasoning is a little more complex for higher copy number plasmids because conjugation can act to change within-host frequencies, rather than entirely replace one plasmid variant. The fundamental intuition, however, remains the same: in a co-infected cell with average within-host variant frequency 1/2, displacement must occur, on average, half as often as co-infection. A similar argument also applies to explaining 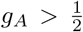 (i.e. the probability of co-conjugation to a singly infected cell resulting in co-infection).

## 4 Supplementary Figures

**Figure S6:**
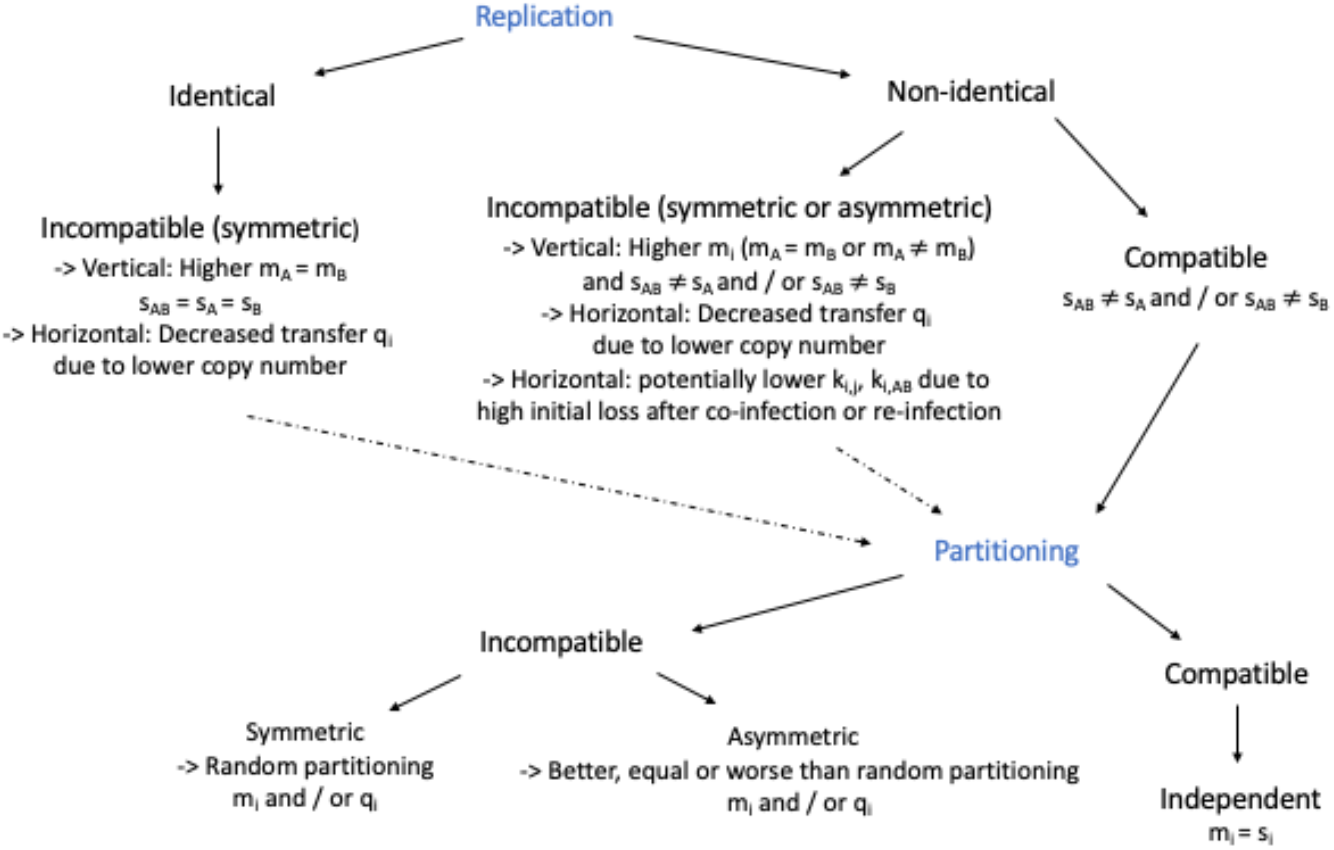
Impact of replication and partitioning on co-infection parameters. Summary of the potential effects of replication and partitioning system relatedness on co-infection modelling parameters as explained in main text section 3.

## 5 Supplementary Tables

**Table S2:** Transmission and fitness effects of biological mechanisms underlying co-infection processes and their empirical values. Here we discuss only direct effects, but all effects on host cell fitness and hence, growth, will affect vertical plasmid transmission indirectly. Similarly, effects from vertical transmission will indirectly affect horizontal transmission through the number of available plasmid donors.

| Biological mechanisms | Effect on vertical plasmid transmission | Effect on horizontal plasmid transmission | Effect on host cell | Empirical values |
| --- | --- | --- | --- | --- |
| Crosstalk in replication regulation | Random replication leads to higher variation in copy number and higher segregation loss of one or both plasmids | None directly | Potentially lower cost through down-regulation of plasmid replication (less than double the replication burden) | 1-15% [4] and 16-22% [5] segregation loss probability per generation for identical plasmids |

| Biological mechanisms | Effect on vertical plasmid transmission | Effect on horizontal plasmid transmission | Effect on host cell fitness | Empirical values |
| --- | --- | --- | --- | --- |
| Crosstalk in partitioning components | Higher segregation loss | None directly | None directly | up to 3% [6]; 5% [7] loss probability per generation |
| Stochasticity in plasmid inheritance | Segregation loss | None directly | None directly | $10^{-3} - 10^{-6}h^{-1}$ [8]; $10^{-3} - 10^{-8}h^{-1}$ [9]; $< 10^{-5}$ for high copy number plasmids [10]; 1-5% of cells in a growing population are empty [11] |
| TA-induced stabilization | Promotes TA-carrying plasmid maintenance and affects non-TA-carrying plasmid inheritance | None directly | Metabolic (potentially temporary) inhibition of plasmid-free cells or TA-plasmid carrying cells | 2.5-100 fold stability increase (single infection) [12]; up to 63% loss of initially resident plasmid in 25 generations of co-infection [13]; more than 95% loss of co-resident plasmid in 100 generations [14] |
| Epistasis in plasmid costs | Unclear; depends on the underlying mechanism | Unclear; depends on the underlying mechanism | Reduction of plasmid burden | Relative fitness in pairwise competition with the wildtype: 0.87 and 0.99 for singly infected and 0.88 for co-infected cells [15] |
| Fertility inhibition systems | None directly | Inhibition of (self and/or co-resident) plasmid transfer | Reduction of plasmid burden through inhibition of conjugation | 100-10,000-fold reduction in conjugation [16] |
| Co-integrates of different plasmid backbones | Unclear | Higher co-transfer and less single transfer; Higher or lower conjugation frequency | Unclear | Average conjugation frequency (i.e. tranconjugants per donor) of single plasmids $8.0 \times 10^{-4}$ and $2.1 \times 10^{-4}$ and co-integrate $3.5 \times 10^{-3}$ in broth [17] |
| Synchronized de-repression of conjugation machinery during co-infection | None directly | Co-transfer of co-resident plasmids | Unclear | Co-transfer is limited by the plasmid with the lower conjugation rate [18] |

